# Modeling the genetic footprint of fluctuating balancing selection: From the local to the genomic scale

**DOI:** 10.1101/2022.07.15.500223

**Authors:** Meike J. Wittmann, Sylvain Mousset, Joachim Hermisson

## Abstract

Natural selection not only affects the actual loci under selection but also leaves “footprints” in patterns of genetic variation in linked genetic regions. This offers exciting opportunities for inferring selection and for understanding the processes shaping levels of genetic variation in natural populations. Here we develop analytical approximations based on coalescent theory to characterize the genetic footprint of a complex, but potentially common type of natural selection: balancing selection with seasonally fluctuating allele frequencies. We show that seasonal allele frequency fluctuations can have important (and partly unexpected) consequences for the genetic footprint of balancing selection. As also confirmed by stochastic simulations, fluctuating balancing selection generally leads to an increase in genetic diversity close to the selected site, the effect of balancing selection, but reduces diversity further away from the selected site, which is a consequence of the allele-frequency fluctuations effectively producing recurrent bottlenecks of allelic backgrounds. This negative effect usually outweighs the positive effect when averaging diversity levels across the entire chromosome. Strong fluctuating balancing selection even induces a loss of genetic variation in unlinked regions, e.g. on different chromosomes. If many loci in the genome are simultaneously under fluctuating balancing selection this could lead to substantial genome-wide reductions in genetic diversity. This may be the case, even if allele-frequency fluctuations are so small that individual footprints are hard to detect. Thus, together with genetic drift, selective sweeps and background selection, fluctuating selection could be one of the major forces shaping levels of genetic diversity in natural populations.

**Article summary:** In some species with multiple generations per year, many loci in the genome experience strong seasonally fluctuating selection, in some cases with stable maintenance of polymorphism. Here we investigate the consequences for levels of genetic diversity at linked neutral sites. Using analytical approximations and stochastic simulations, we find a characteristic local genetic footprint with a diversity peak around the selected site and a diversity valley further away and a substantial reduction in diversity levels chromosome-wide and even genome-wide.

## Introduction

One of the key insights of modern population genetics is that natural selection at one locus in the genome can influence patterns of genetic variation at other loci, in particular at closely linked sites (Barton 2000; Cutter and Payseur 2013; Maynard Smith and Haigh 1974). Characterizing these genetic footprints of selection is important for two main reasons. Expected patterns of genetic diversity derived from models can be used to scan genomes for regions under selection or to infer the type of selection acting in a candidate region (Gillespie 1997; Vitti *et al*. 2013). Moreover, cumulatively, the effects of linked selection at many loci contribute to shaping levels of genome-wide variation at neutral loci but also other selected loci (Corbett-Detig *et al*. 2015; Huang *et al*. 2014; Leffler *et al*. 2012; Sella *et al*. 2009). Ultimately, genetic footprints therefore also affect a population’s ability to adapt to a changing environment.

The genetic footprints of positive selection (hard and soft selective sweeps) and purifying selection (background selection) have been characterized in detail in numerous studies (for recent reviews see Charlesworth and Jensen 2021; Hermisson and Pennings 2017). Likewise, footprints of balancing selection, where two alleles are maintained at (roughly) constant frequency, are well-understood (e.g. Charlesworth 2006; Fijarczyk and Babik 2015; Gao *et al*. 2015). However, other typical modes of selection in natural populations have received less attention. One of them is temporally fluctuating selection. Many natural populations experience bouts of strong selection with frequent, more or less regular changes in the direction of selection (Bell 2010; Moorcroft *et al*. 1996; Nicolaus *et al*. 2016; Siepielski *et al*. 2009; Thurman and Barrett 2016). At the genetic level, such fluctuating selection can cause rapid changes in allele frequencies over short time scales (Abdul-Rahman *et al*. 2021; Garcia-Elfring *et al*. 2021; Rudman *et al*. 2022). For example, Bergland *et al*. (2014) found large seasonal allele-frequency fluctuations at hundreds of loci across the genome in a temperate *Drosophila, melanogaster* population. Many of these seasonal SNPs shift in parallel across multiple European and American populations and are also consistent with clinal variation (Machado *et al*. 2021). Recently, large allele-frequency fluctuations due to fluctuating environmental conditions have also been found in the non-biting midge *Chironomus riparius* (Pfenninger and Foucault 2022; Pfenninger *et al*. 2022).

Persistent fluctuating selection at a locus can only be observed if polymorphism is maintained. Several plausible mechanisms for the long-term maintenance of genetic variation due to selection exist. In particular, fluctuating selection itself can maintain genetic polymorphism under appropriate conditions (e.g. Gillespie 1973, 1978; Haldane and Jayakar 1963; Park and Kim 2019; Wittmann *et al*. 2017; Yi and Dean 2013). Alternatively, it could act in conjunction with other forces maintaining variation, such as negative frequency-dependent selection. In Bergland et al.’s *Drosophila* study, many of the fluctuating polymorphisms found appear to be long-term stable and are shared with other *Drosophila* populations around the world or even with the sister species *Drosophila simulans*. Also in *Chironomus riparius*, some of the loci under fluctuating selection exhibit signals of long-term balancing selection (Pfenninger *et al*. 2022). In this paper, we focus on this scenario of “fluctuating balancing selection”, where the alleles of a stable polymorphism exhibit temporal fluctuations and explore the corresponding genetic footprint.

Since fluctuating balancing selection is essentially a combination of classical balancing selection maintaining a polymorphism and strong directional selection, leading to rapid changes in allele frequencies, well-known results for these limiting scenarios can give us first clues to its footprint. Classical balancing selection leads to an increase in diversity at closely linked neutral sites (Charlesworth 2006). This is because close to the selected site, where the recombination rate between neutral and selected locus is small, there is little exchange between the two allelic “subpopulations”. They become differentiated from each other, and samples including chromosomes from different subpopulations harbor substantial diversity. By contrast, rapid changes in allele frequencies, as in a selective sweep, tend to reduce genetic diversity at linked neutral sites (Maynard Smith and Haigh 1974; Stephan 2019), where alleles can hitchhike to high frequencies or to fixation. More generally, diversity is reduced by hitchhiking when effects of selection in the genomic background lead to an increased offspring number variance at the focal locus (Barton 2000). This also holds if the strength and direction of linked selection fluctuates with time.

Numerical results by Gillespie (1997) and Taylor (2013) for uncorrelated random environmental fluctuations and by Park and Kim (2019) for a model with cyclical selection suggest that fluctuating balancing selection can both increase and decrease diversity at linked neutral sites. However, there is a lack of analytical results and the case of cyclically fluctuating selection has not been explored in depth. Here, we consider strong seasonal selection, where allele frequencies fluctuate on time-scales much faster than genetic drift. We derive analytical results to investigate how the genome-wide footprint depends on the strength and dominance of selection and the fluctuation period. We discuss whether and when the positive or negative effects of fluctuating balancing selection on diversity levels dominate on the scale of chromosomes and genomes.

## Model and Methods

### Selection model and allele-frequency trajectories

As our standard model, we consider a diploid population of size *N* living in a seasonal environment. Generations are discrete, and every year there are *g_w_* generations of winter followed by *g_s_* generations of summer, with *g_total_* = *g_w_* + *g_s_*. We initially assume that selection acts at a single locus with two alleles, which we will call the *winter allele W*, and the *summer allele S*. We assume that selection follows a yearly cycle such that the fitness of a genotype (WW, WS, or SS) depends on the generation of the year *i* = *t* mod *g_total_* (Table 1), where *t* is time in generations and mod is the modulo operator. The winter allele is favored in winter with selection coefficient *s_w_* and dominance coefficient *h_w_*, and the summer allele in summer with selection coefficient *s_s_* and dominance coefficient *h_s_*. While we focus on symmetric scenarios with *s_w_* = *s_s_* = *s, h_w_* = *h_s_* = *h*, and *g_w_* = *g_s_* = *g*, we will also explore the role of asymmetry. In addition, we will later relax the assumption of cyclical selection to consider random switches between seasons.

**Table 1.**
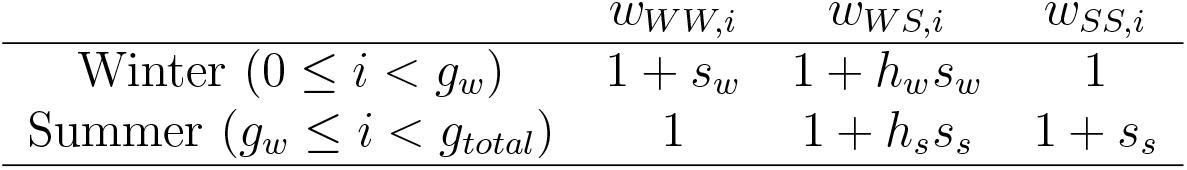
Fitness values *w_WW,i_, W_WS,i_*, and *w_SS,i_* for the three genotypes in winter and summer. The index *i* denotes the generation of the year (starting at 0 for the first winter generation).

Stable polymorphism in this model occurs if there is marginal overdominance (Haldane and Jayakar 1963), that is, if heterozygotes have a higher geometric mean fitness than either homozygote, so that each allele can invade when rare. If *s_w_* = *s_s_* = *s* and *h_w_* = *h_s_* = h, polymorphism is stable if (1 + *hs*)^2^ > 1 + *s* or equivalently 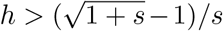. In other words, the currently beneficial allele needs to be sufficiently dominant. Note that *h* = 0.5 fulfills this condition. If *h* ≠ 0.5, dominance switches between seasons. Coexistence in this model is a special case of the “segregation lift” mechanism for the maintenance of multi-locus polymorphism under cyclically fluctuating selection (Wittmann *et al*. 2017). As mentioned above, multiple other mechanisms also give rise to fluctuating balancing selection. For example, seasonally fluctuating selection coefficients could act in conjunction with rare-type advantage, e.g. due to specialization on different resources or differences in the interactions with other species. As discussed below, most of our results do not depend on the specific selection model, but only on the allele-frequency trajectories.

We assume the following life cycle: Each generation t starts with an infinite gamete pool, at which point also the allele frequency *p_t_* is quantified. Second, symmetric mutation occurs with probability *u* per allele copy per generation. Third, *N* diploid zygotes form by random union of gametes to give rise to the adult population. Finally, adults contribute to the gamete pool of the next generation in proportion to their fitness values. If we let the population size go to infinity, the dynamics become deterministic, and we can compute the allele frequency of the winter allele in generation *t* + 1, *p*_*t*+1_, from the allele frequency in generation *t*, *p_t_*, in two steps accounting for mutation,

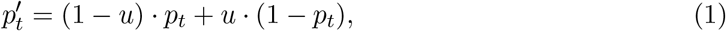

and selection,

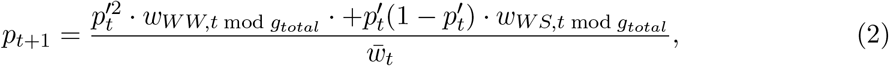

where 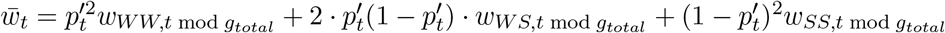 is the mean fitness of diploid adults in generation *t*. Note that with random union of gametes, the population is in Hardy-Weinberg equilibrium before selection.

In this study, we assume that the same selection dynamics have been acting for a long time. We thus need to characterize the long-term behavior of (1) and (2). In situations where polymorphism is maintained, we obtain, independently of the initial conditions, a cyclical allele-frequency trajectory described by winter-allele frequencies *π_i_* for 0 ≤ *i* < *g_total_* such that 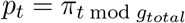. In symmetric cases, we show in Appendix 1 that

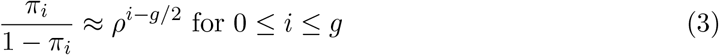

with 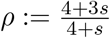, and thus

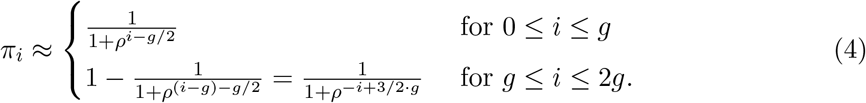

Note that *π*_0_ = *π*_2*g*_ and that both expressions coincide for *i* = *g*.

We find excellent agreement between this analytic approach and stochastic simulations with *N* = 10,000 (Fig. 1, see below for details of the stochastic simulation methods). Relevant summary statistics of the allele-frequency trajectories are shown in Fig. 2. The magnitude of fluctuations increases with both the number of generations per season (Figs. 1 and 2 A) and with the selection coefficient (Fig. 2 B), but is barely affected by the dominance coefficient and the mutation rate (Fig. 2 C, D).

**Figure 1.**
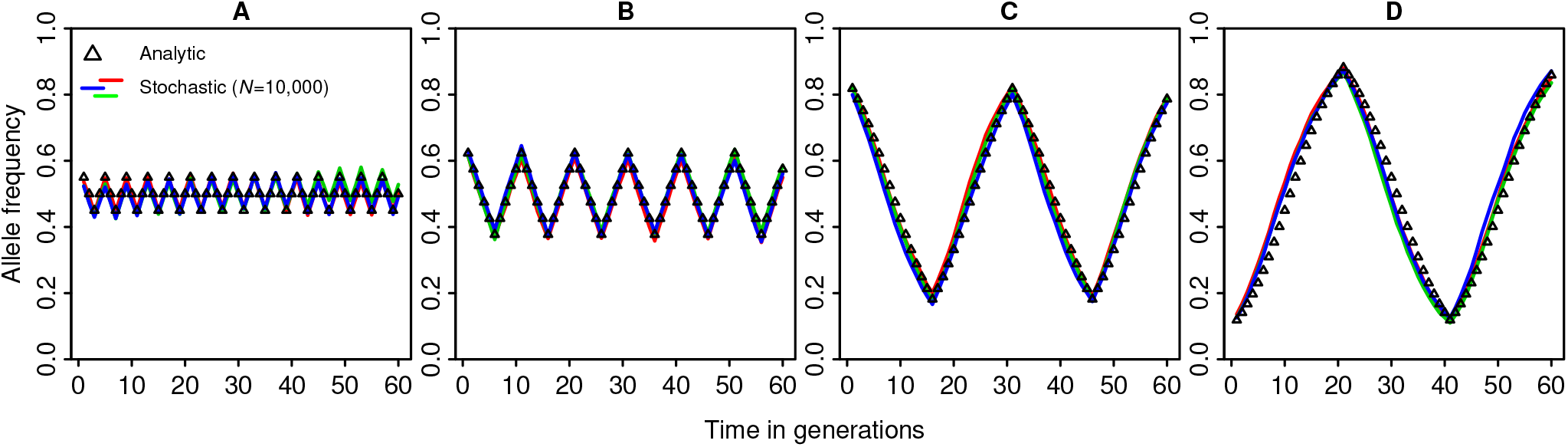
Comparison of deterministic allele-frequency trajectories as determined analytically using (4) with three replicate stochastic simulations (conducted via the Python package simuPOP, see below for details) shown in different colors, though overlapping to a large extent. The panels differ in season length, *g*. A) *g* = 2, B) *g* = 5, C) *g* = 15, and D) *g* = 20. Other parameters: *s* = 0.5, *h* = 0.6, *u* = 10^-6^.

**Figure 2.**
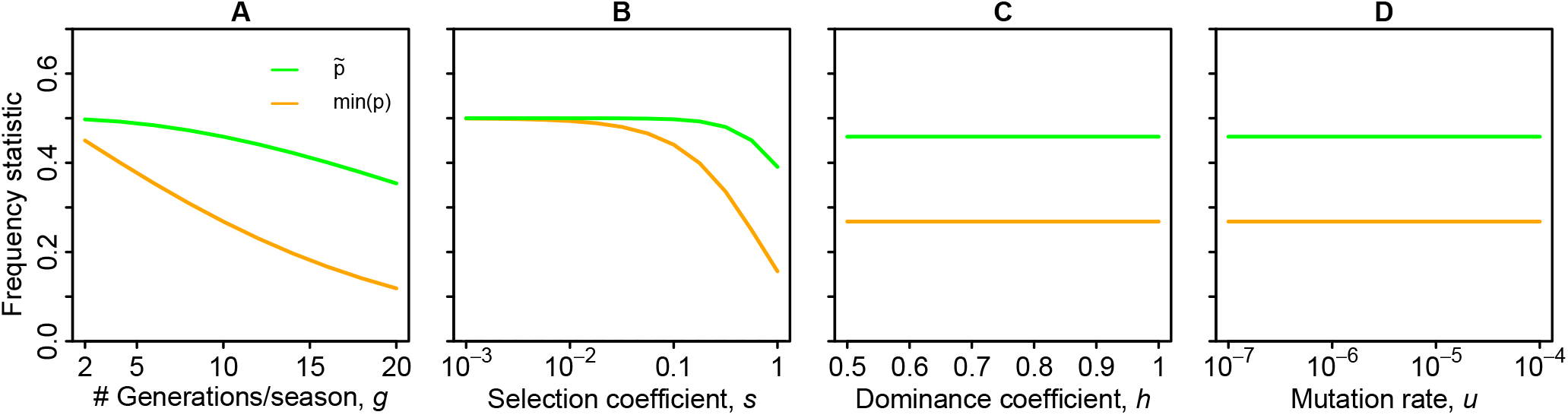
Statistics of cyclical allele frequency trajectories calculated using the analytic approximation in (4) (lines). 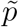 is the harmonic mean allele frequency that is important for the quantification of diversity below. The minimum winter allele frequency in a cycle, min(*p*), serves as a measure for the magnitude of fluctuations. Other parameters: *N* = 10, 000, *u* = 10^-6^, *g* = 10, *s* = 0.5, *h* = 0.6.

Although we focus mainly on seasonal (periodic) fluctuations, we also present some results for an uncorrelated random environment, where selection in each generation is of type W or S independently and with equal probability in every generation (see Table 1). Since autocorrelation is irrelevant for the stability of polymorphism by marginal overdominance (Gillespie 1973), the conditions for stable polymorphism are the same as in the seasonal model.

### Quantifying expected diversity levels

We now turn our attention to the linked variation at either side of the selected locus, and consider a biallelic neutral locus at recombination distance *r* (with recombination probability 0 ≤ *r* ≤ 0.5). Symmetric mutation at this locus changes its allelic state with probability *u* per generation. Our quantity of interest is the expected heterozygosity, defined as the probability that two randomly chosen chromosomes have different alleles at the site of interest. With symmetric biallelic mutation, this requires an odd number of mutations on the ancestral lineages connecting the two sampled chromosomes with their common ancestor.

Fluctuating balancing selection affects linked diversity in two main ways, which we will address in two separate steps in our analytical approximation. First, balancing selection (fluctuating or not) that maintains two alleles at the selected locus for a long time, structures the population into two parts according to the allelic state: winter and summer. Ancestral lines of descent from different parts of this genetically structured population can only coalesce after a recombination event along these lines unites them in the same background. This effect is most conveniently described by a structured coalescent model. Second, rapid allele frequency changes due to fluctuating selection increase the offspring variance not only at the selected site, but also at linked (and even unlinked) neutral sites due to the hitchhiking effect (Hill and Robertson 1966; Maynard Smith and Haigh 1974). Following Barton (2000), this effect can be captured by an appropriate rescaling of the effective population size.

### Analytical approximation step 1: structured coalescent

In the first step of our approximation, we use a version of the classical structured coalescent model (Hudson and Kaplan 1988; Kaplan *et al*. 1988) to derive the expected time to the most recent common ancestor (coalescence) for samples of size two from a neutral locus at recombination distance *r* from the selected site. Each lineage at the neutral locus can be either in a winter or summer background, meaning that the chromosome carries the winter or the summer allele, respectively, at the selected locus. We trace the ancestry of two sampled lineages backward in time and keep track of their genetic background. Formally, for a sample of size two, this leads to a Markov chain with four states: state (2,0) with both lineages in the *W* (winter allele) background, state (0,2) with both lineages in the *S* (summer allele) background, state (1,1) with one lineage in each background, and finally the absorbing state (*) where both lineages have coalesced. Coalescence is only possible if both lineages are in the same background. Lineages can switch to the other background either due to a recombination between neutral and selected site or due to a mutation at the selected site.

Given an allele frequency *p* for a focal allele at the selected locus, lineages within this background coalesce at rate 1/(2*Np*), they recombine into the other background with rate *r*(1 – *p*) and they mutate into the other background with rate *u*(1 – *p*)/*p*. In the standard model of balancing selection, allele frequencies are constant over time. In our model, however, allele frequencies and thus transition probabilities vary cyclically over time. This would make our Markov chain time-inhomogeneous and analytically intractable. In the first step of our approximation, we therefore assume that *N* is sufficiently large and that *u* and *r* are small such that coalescence, mutation, and recombination probabilities are small per generation and even per year, and the probability of multiple events per year is negligible (we relax this assumption in the second step of our approximation below). We can then approximate the ancestral process by a homogeneous continuous-time Markov process whose rates are given by the respective average transition probabilities over the cyclic allele-frequency trajectory as given by the frequencies *π_i_*. The coalescence rate per generation in the new process is

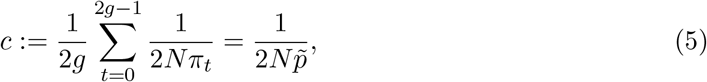

where 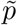 is the harmonic mean allele frequency over the cycle. Here we focus on symmetric situations, such that both alleles have the same harmonic mean frequency, i.e. the harmonic mean of *π_t_* is equal to the harmonic mean of 1 – *π_t_* (see Appendix 6 for asymmetric cases). Next, we consider the transition rate between backgrounds due to mutation:

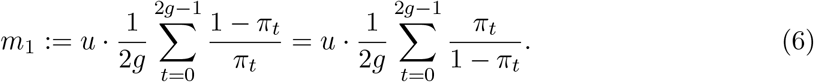

Again, the second equality holds because of symmetry, so that the transition rate is the same in both directions. *m*_1_ can be expressed in terms of the harmonic mean:

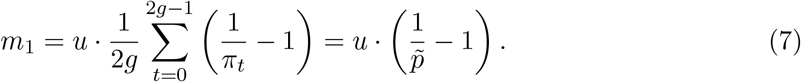

Finally, we consider the transition rate between backgrounds due to recombination:

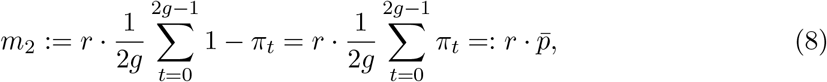

where 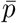 is the arithmetic mean frequency along the cycle, i.e. 0.5 in symmetric cases. Let *m* = *m*_1_ + *m*_2_ be the total rate at which lineages switch to the respective other background due to mutation or recombination.

We now use these transition rates to compute the expected time to coalescence for a sample of two lineages via first-step analysis:

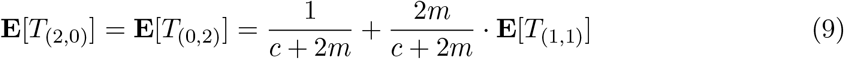

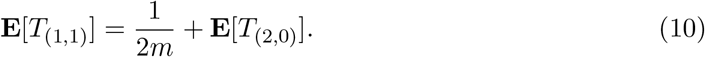

Solving the recursion, we obtain

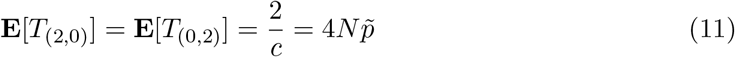

and

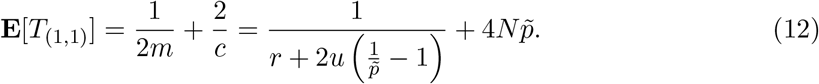

For symmetric selection scenarios our analytical approximation only depends on population size, mutation rate, recombination rate, and the harmonic mean allele frequency over the cycle, which can be computed from the analytical approximation for the cyclical allelefrequency trajectory in (4) as

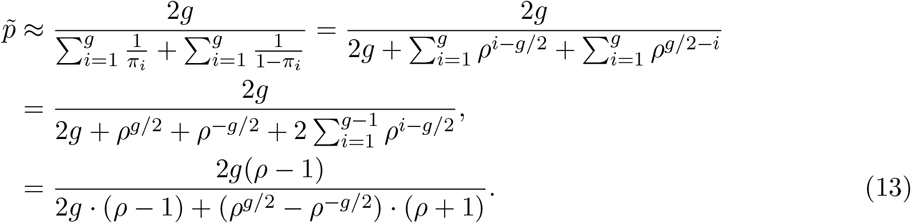

Note that the harmonic mean allele frequency does not depend on the ordering of allele frequencies in a cycle and can take the same value also for cases with different cycle lengths.

Finally, given the expected coalescence time for sample configuration (*i,j*), we approximate the corresponding expected heterozygosity as

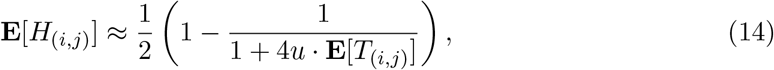

which accounts for back-mutation (derivation in Appendix 2). Since sampling in our model happens in the gamete pool, i.e. before selection, we can assume Hardy-Weinberg equilibrium, and the expected heterozygosity in a random sample at time *t* is

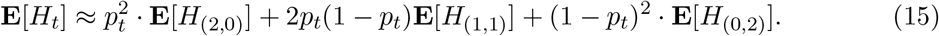

We measure the effect of linked selection by comparing **E**[*H_t_*] to the neutral baseline, which is obtained by substituting the expected neutral coalescence time **E**[*T*_(*i,j*)_] = 2*N* in (14).

If the mutation rate is small relative to recombination, we can further approximate

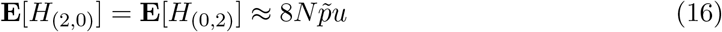

and

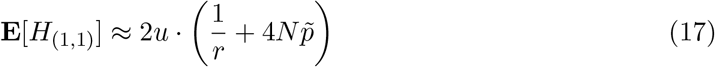

to leading order in *u*, and thus

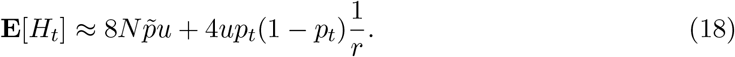

The % change relative to the neutral expectation (≈ 4*Nu*),

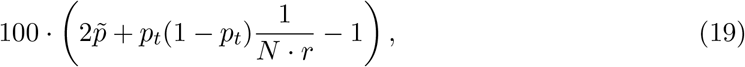

then only depends on the compound parameter *N* · *r*. The underlying biological reason is that the decrease of the diversity close to the selected site only depends on the relative size of the two time scales for coalescence (~ *N*) and recombination ~ (1/*r*). With large *N* · *r*, recombination is fast enough that it mixes backgrounds before coalescence typically occurs, and the genetic structure does not play a large role anymore.

Equation (15) (or (18) for small mutation rate *u* ≪ *r*) complete the first step of our analytical approximation, which is expected to be accurate as long as recombination is rare on the timescale of the fluctuation period (or season).

### Analytic approximation step 2: Hitchhiking

In our approximation so far, we have assumed that all ancestral lineages experience an averaged environment of summer and winter generations on their course back in time to the first non-trivial genealogical event, with can be mutation, recombination, or coalescence. For seasonal fluctuations and realistic population sizes and mutation rates, this is generally true for mutation and coalescence. With loose linkage and recombination probabilities up to 0.5, however, lineages will typically switch backgrounds multiple times per cycle. On these short timescales, random associations of lineages at the neutral locus with either the *W* -or *S*-allele do not average out, but contribute to an increased offspring-number variance among lineages, corresponding to a decreased variance-effective population size.

The second step of our approximation accounts for this hitchhiking effect of fluctuating selection with loose linkage. Instead of extending the structured coalescent to a time-inhomogeneous Markov process, we use a method by Barton (2000) to derive the effect of the fluctuations. To this end, we start from Eq. (A6) in Barton (2000), which states that the hitchhiking effect of fluctuating selection decreases the variance-effective population sizes by a factor *F*, i.e. *_e_* = *N/F*, with

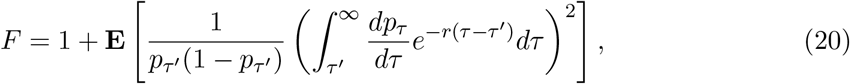

where the expectation is over the time course of allele frequencies 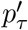. Note that we use *τ*’ as lower integration boundary, because the 0 in Barton (2000) appears to be a typo. Discretizing this equation to match our discrete-time model leads to

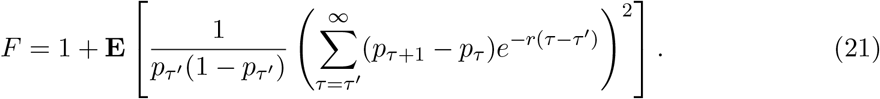

For the uncorrelated selection scenario we use this equation directly, but with an upper summation boundary of *τ*’ + 500. For cyclical selection, we can further simplify:

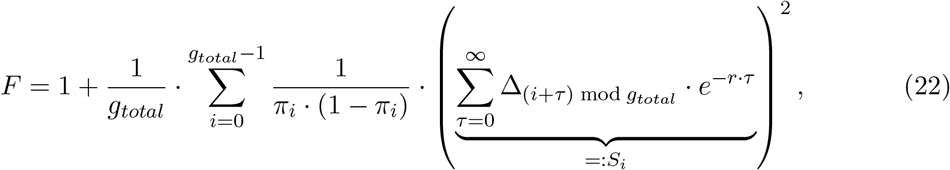

where

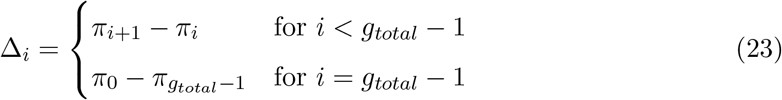

are the allele-frequency increments from generation to generation in the cycle.

To evaluate the infinite sum, we split up the terms according to position in the cycle and then use one geometric series per point in the cycle:

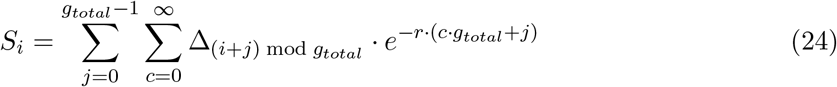

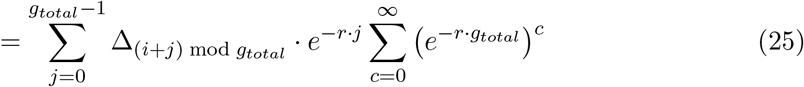

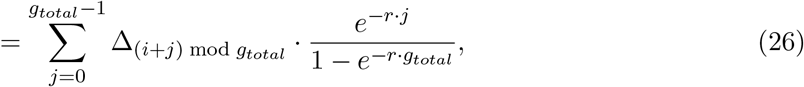

which completes our expression for *F*.

To see how this expression can be combined with the 1st step of our analytical approximation, note that we can also compute an effective population size in the structured coalescent model if we assume that *u* << *r*, and 1/*N* << *r* << 1/*g*. That is, recombination within a cycle is rare, but recombination is much faster than coalescence so that lineages switch between backgrounds often enough that they reach a stationary distribution. In this stationary distribution, the probability to be in a certain background at a certain time is just the frequency of the respective background. In the more general (potentially asymmetric case), the coalescence rate is then

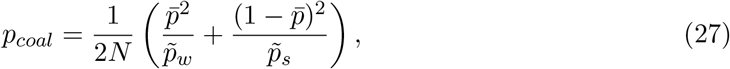

where 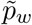 and 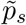 are the harmonic mean allele frequencies of the two alleles. This expression reduces to 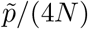 in the symmetric case. We can then compute 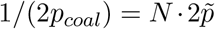. Via this effective population size, we can now connect the factor F which also takes into account possible background switches within a cycle to the structured coalescent approximation. That is, we complete our analytical approximation by substituting 1/(2F) for 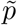 in (11) and (12).

In Appendix 3 we verify that *F* converges to 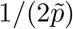 as the recombination rate between neutral and selected locus approaches zero. Thus, for small recombination distances, this extended approximation reduces to the one from our first step, but it captures the phenomenon that lineages can switch background multiple times per season for large recombination distances. Also note that unlike in the first analytical approximation, the ordering of allele-frequencies in a cycle and the cycle length can now have an effect via *F*.

### Stochastic individual-based simulations

To check our analytical results, we ran individual-based simulations using the Python library simuPOP version 1.1.7 (Peng and Kimmel 2005). Because of computational constraints, we did not simulate entire chromosomes of several millions to billions of sites. Instead, we simulated *L* sites (e.g. 40,000) linked to the selected locus and placed them strategically, so that we obtain more data close to the selected site where patterns of variation are expected to change on a smaller scale than far away from the selected locus. Specifically, we set a minimum recombination probability *r_min_* = 10^-8^ and a maximum recombination probability *r_max_* = 0.49 with the selected site and placed the ith simulated site at a recombination distance of 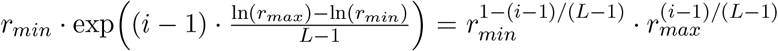 from the selected site. That is, the sites were evenly spaced on a logarithmic scale. To explore the genomewide effects of multiple loci under fluctuating balancing selection, we will also extend our simulation approach to multiple loci under selection, as described below.

To initialize a simulation run, we drew from a simulated distribution of allele counts under neutrality (see Appendix 4 for details). This was also done for the selected loci. That is, the start of the simulation can also be seen as the onset of selection. Unless stated otherwise, all simulations ran for 15 · *N* generations. Samples of 1,000 individuals (or the entire population if *N* ≤ 1, 000) were taken roughly every 0.5 · N generations, with the precise sampling interval fine-tuned such that sampling always happens after the first generation of a year. Unless stated otherwise, we only used the last five sampling points (at roughly 13*N*, 13.5*N*,..., 15*N* generations) as a basis for our results. Our measure of diversity at site *i* is the heterozygosity 2 · *p_i_* · (1 – *p_i_*), where *p_i_* is the sample allele frequency at site *i*. Heterozygosities at all 40,000 linked sites (i.e. excluding the selected sites) were put into 25 bins of equal size (i.e. the 1600 sites closest to the selected site, the next closest 1,600 sites), and then averaged. The average change in heterozygosity relative to the neutral expectation is plotted as a function of recombination distance, whose summary value for the respective bin is obtained by averaging at the logarithmic scale and then taking the exponential.

## Results

### Footprint of fluctuating balancing selection

As shown in Fig. 3 for the allele-frequency trajectories in Fig. 1, cyclic fluctuations can have a strong effect on the footprint of balancing selection. While weak fluctuations basically reproduce the well-known footprint of constant balancing selection, with a strongly localized increase in diversity close to the selected site (Fig. 3 A), the effect on diversity becomes nonmonotonic (increasingly U-shaped) along the chromosome for larger fluctuation amplitudes (Fig. 3 B-D).

**Figure 3.**
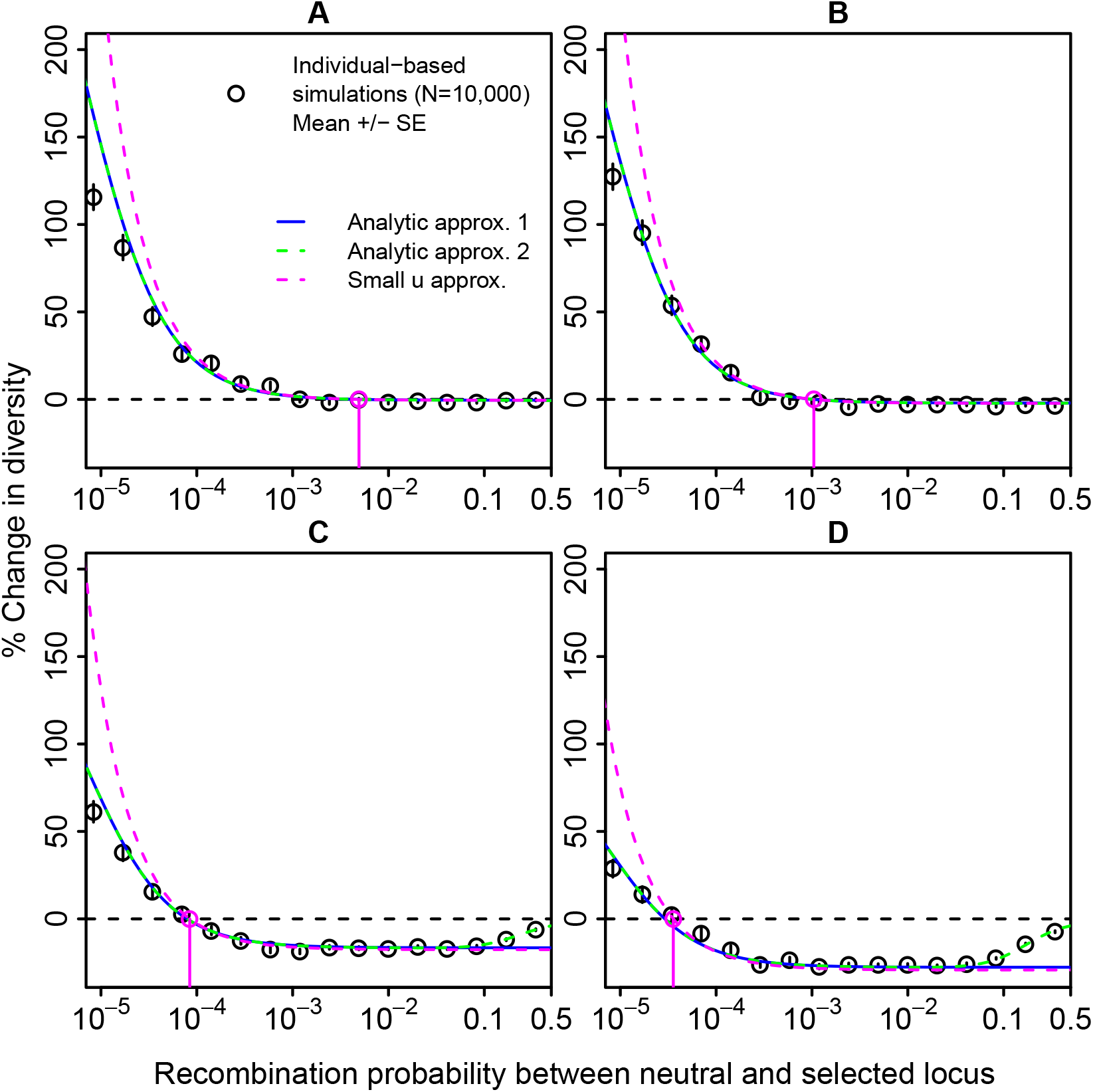
Genetic footprints of fluctuating balancing selection for various season lengths, *g*. A) *g* = 2, B) *g* = 5, C) *g* = 15, and D) *g* = 20. The y-axis represents the relative change in diversity compared to the neutral case. Points are averages over 100 replicates with five sampling points each. See Fig. 1 for the corresponding allele-frequency trajectories. The pink vertical line and circle indicate the recombination distance at which expected heterozygosity falls below neutral levels as estimated by (28). A zoom into the region of large recombination rates is shown in Fig. A2 and an extension to lower recombination rates is shown in Fig. A3.

In the limit of very small fluctuations around 0.5, we have 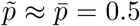 and our approximation for the coalescence times (11) reduces to **E**[T(_2_,_0_)] = **E**[*T*_(0,2)_] = 2*N*, which equals the expected coalescence time under neutrality, and **E**[*T*_(1;1)_] > 2*N*. Thus, the expected heterozygosity in a random sample from the population is always larger than under neutrality, but approaches the neutral baseline with increasing recombination distance from the selected site (see (12)). For *r* → 0, the equilibrium diversity is only limited by allelic turnover due to recurrent mutation at the selected site and can get very large (%-change ~ 1/(*uN*)). Note that our simulations predict slightly lower diversity levels in this regions (see also Fig. A3). The reason appears to be that, due to computational limitations, we could not run the stochastic simulations long enough for very high levels of diversity to build up (see also Fig. A3 and more detailed discussion in Appendix 5).

As allele-frequency fluctuations increase in magnitude (for example with increasing number of generations per season in Fig. 3 B, C, D), the characteristic footprint of fluctuating balancing selection emerges. While close to the selected site, diversity is still increased, it declines below the neutral baseline with increasing recombination distance from the selected site, before it partially recovers at even larger distances. The mathematical cause for this diversity valley can be observed from equations (11) and (12). For fluctuating frequencies, the harmonic mean, 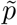, drops below the arithmetic mean, 0.5, reducing the expected coalescence times. Biologically, we can distinguish two effects of the fluctuations that are both mediated by the harmonic mean. First, both allelic backgrounds experience regular bottlenecks, i.e. times of low frequency, during which pairs of lineages in these background have an increased coalescence rate. This first effect can also be interpreted as a reduction of the effective population size. Second, transitions between backgrounds due to recurrent mutation at the selected locus (allelic turnover) increase with increasing magnitude of fluctuations and thus decreasing 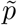 (12). While the first effect determines the bottom of the diversity valley, the second effect (only) contributes to a faster decline towards this minimum.

When the recombination distance to the selected locus increases further and approaches 0.5, diversity levels leave the “valley” and start to recover (see Fig. 3 C,D). This effect is not captured by the first step of our approximation, but only by the second step, which accounts for the fact that with larger *r* lineages are increasingly able to “track” allele frequencies through multiple background switches per year. Lineages can then escape a shrinking background quickly enough to not be forced together through a bottleneck, where coalescence is most likely. Strong recombination thus “reduces the reduction” of the effective population size due to the fluctuations. However, diversity levels do not fully return to neutral levels even for free recombination with *r* = 0.5 (see Fig. 3 D). The consequences of this will be explored below when we look at the genome-wide effect of fluctuating balancing selection.

#### Quantifying the diversity valley

We can use our analytical approximations to quantify the hallmarks of the diversity valley. From (18), the recombination rate *r*\* where heterozygosity levels fall below the neutral baseline results as

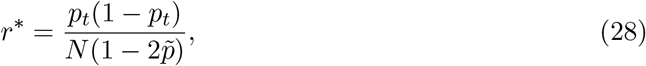

which is shown as a pink vertical line and circle in Fig. 3. Note that, since typically *r*\* » *u*, the weak-mutation approximation can be used to derive this threshold. With increasing magnitude of the fluctuations (and decreasing 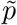), the intersection point moves closer to the selected locus. Since *r*\* ~ 1/*N*, it declines with *N*. In other words, the intersection occurs at a constant value of the compound parameter *rN* for a given level of the fluctuations (fixed 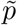). For *r* > *r*\*, the bottom of the valley is reached when the first term in the coalescence-time approximation (12) for samples from different backgrounds can be ignored relative to the second one 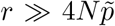. The minimal diversity level is well predicted by all analytic approximations and is given by

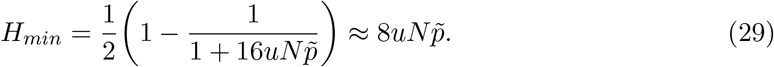

The corresponding percentage change of 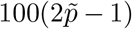 relative to neutral diversity depends on the harmonic mean as the only summary statistic to capture the effect of the fluctuations.

Finally, we observe from Eq. (22) that diversity levels increase above the valley bottom once *r* · *g_total_* is no longer small and recombination during a single selection cycle becomes likely. We thus have a different scaling for both edges of the diversity valley, with the lower end *r*\* ~ 1/*N* and (via 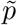) depending on the amplitude of the fluctuations, while the upper end, *r*\*\* ~ 1/*g_total_*, scales with the fluctuation frequency, independently of *N* and the fluctuation amplitude. Since typically 1/*N* ≪ 1/*g_total_* for seasonal fluctuations, the valley can extend over several orders of magnitude of the recombination distance *r*.

Due to the allele-frequency fluctuations, the expected footprint of a random sample also depends on the time in the seasonal cycle in which the sample is taken (Fig. A8). With more balanced allele frequencies in the middle of the season, it is more likely that two lines from different backgrounds end up in a random sample of size 2 and thus there is a stronger increase in diversity close to the selected site. However, the long-range effects are not noticeably affected by the seasonal sampling point.

So far, we have focused on diversity levels long after the onset of cyclical selection pressures. The temporal emergence of these footprints after the onset of fluctuating balancing selection is shown in Fig. A9. The negative effect on diversity develops first, especially close to the selected site. Over time, backgrounds become differentiated and diversity close to the selected site builds up and starts to exceed neutral levels. At the same time, also the diversity reduction further away from the selected site intensifies and approaches the final levels.

#### Effects of model parameters

So far we have focused on how season length affects the genetic footprint via its effect on the magnitude and period of fluctuations. Results for how the other model parameters shape the footprint of fluctuating balancing selection are shown in A4-A7 and all parameter effects are summarized in Fig. 4. The main effect of the mutation rate *u* is to increase the rate of allelic turnover at the selected locus which limits the diversity very close to the selected site. At very large mutation rates, there is additionally a saturation effect (see Fig. A6 and Discussion in Appendix 5). Increasing selection strength *s* increases the amplitude of fluctuations and, like increasing *g*, weakens the diversity increase close to the selected site and deepens the diversity valley. Beyond that, there is a special effect when Ns is small, that is when selection is weak relative to genetic drift. Here the individual-based simulations (Figs. A4 and A5) predict that diversity levels are close to neutral expectations. Since our analytical approaches are based on the assumption of deterministically cycling allele frequencies, they require that selection be strong relative to genetic drift and thus perform poorly for small *s* and small *N*.

**Figure 4.**
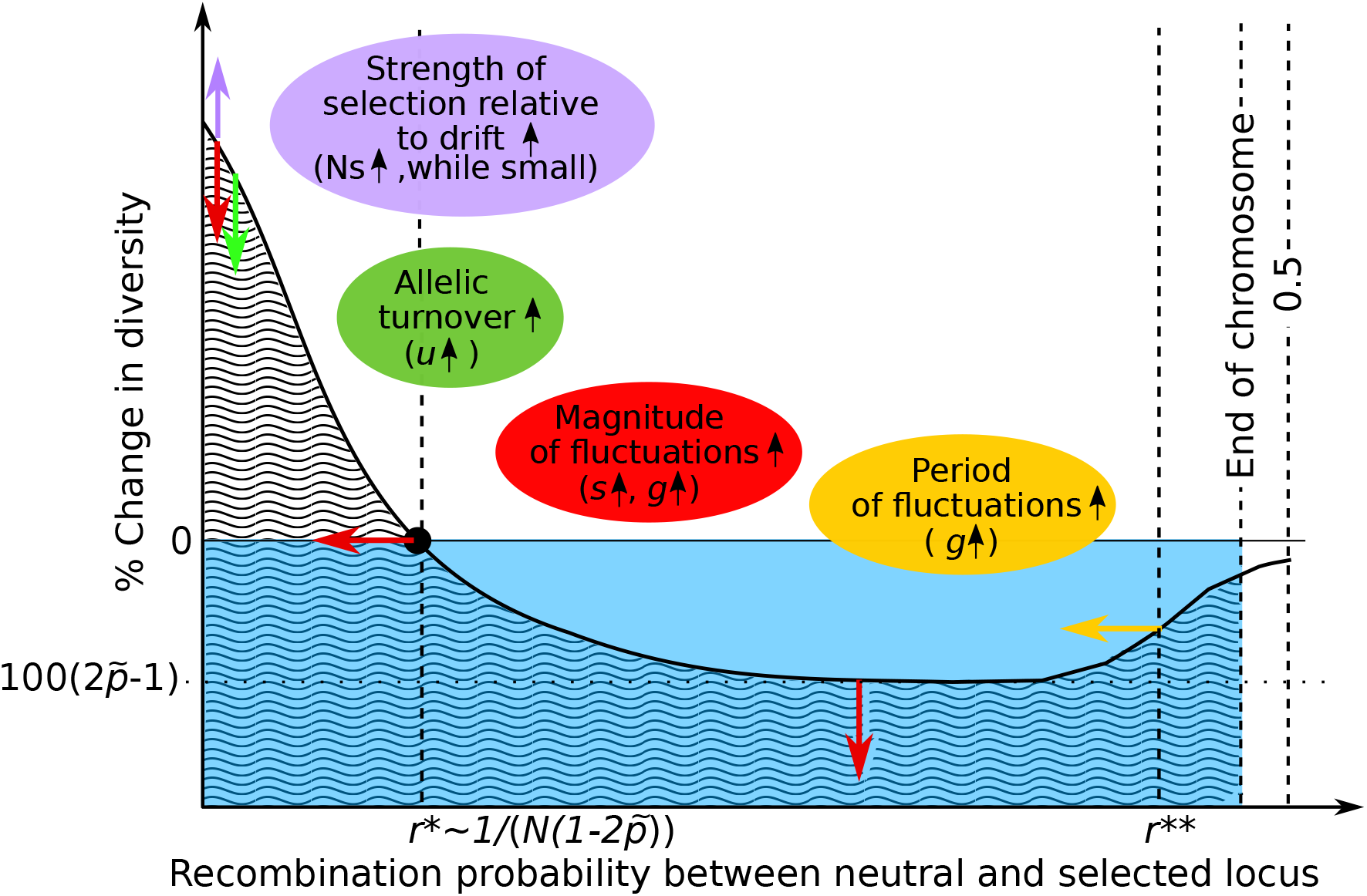
Cartoon illustrating how the various model parameters influence the shape of the genetic footprint of fluctuating balancing selection (area with wave pattern). The blue area corresponds to neutral diversity levels. The arrows indicate the direction in which the shape of the genetic footprint will change when the respective model parameters are increased. The effect of *h* is minimal and not shown. Recall, however, that all of our analyses are based on the assumption that the dominance coefficient is large enough for polymorphism to be long-term stable.

### Asymmetric and random fluctuations

While we focus on symmetric scenarios, we also simulated asymmetric scenarios and extended both analytical approximations accordingly (see Appendix 6). Again, our second analytical approximation provides an excellent fit to the simulations. The results suggest that with increasing asymmetry, the reduction in diversity away from the selected site becomes weaker (Fig. A10). In asymmetric scenarios, there is a substantial reduction in diversity for samples from the rare background close to the selected site, whereas samples from the overall more common background exhibit either a slight increase or a slight decrease in diversity.

A model version with random switches between selection parameters and thus irregular allele frequency fluctuations gives a qualitatively similar genetic footprint as the model with seasonally fluctuating selection (Fig. 5). Again, we compare the results from individual-based simulations to two approximations. However, in this case, the approximations are only semi-analytic because we based them on allele-frequency trajectories obtained from iterating the deterministic model equations (1) and (2) for 100,000 generations with random environments. From these trajectories, we then calculated the frequency statistics for (11) and (12). We also used the trajectories to evaluate (21). Once again, the first step of the approximation predicts the minimum diversity levels along the chromosome, but not the recovery at high recombination distances. The second step of the approximation also captures this recovery. Compared to our results for the seasonal selection model, diversity levels recover from this valley bottom at smaller recombination rates. The likely reason is that the largest fluctuation amplitudes in the random-selection model typically occur in much longer excursions of the allele-frequency trajectory above or below the arithmetic mean than in a yearly cycle, thus giving lineages more time to escape from the smaller background.

**Figure 5.**
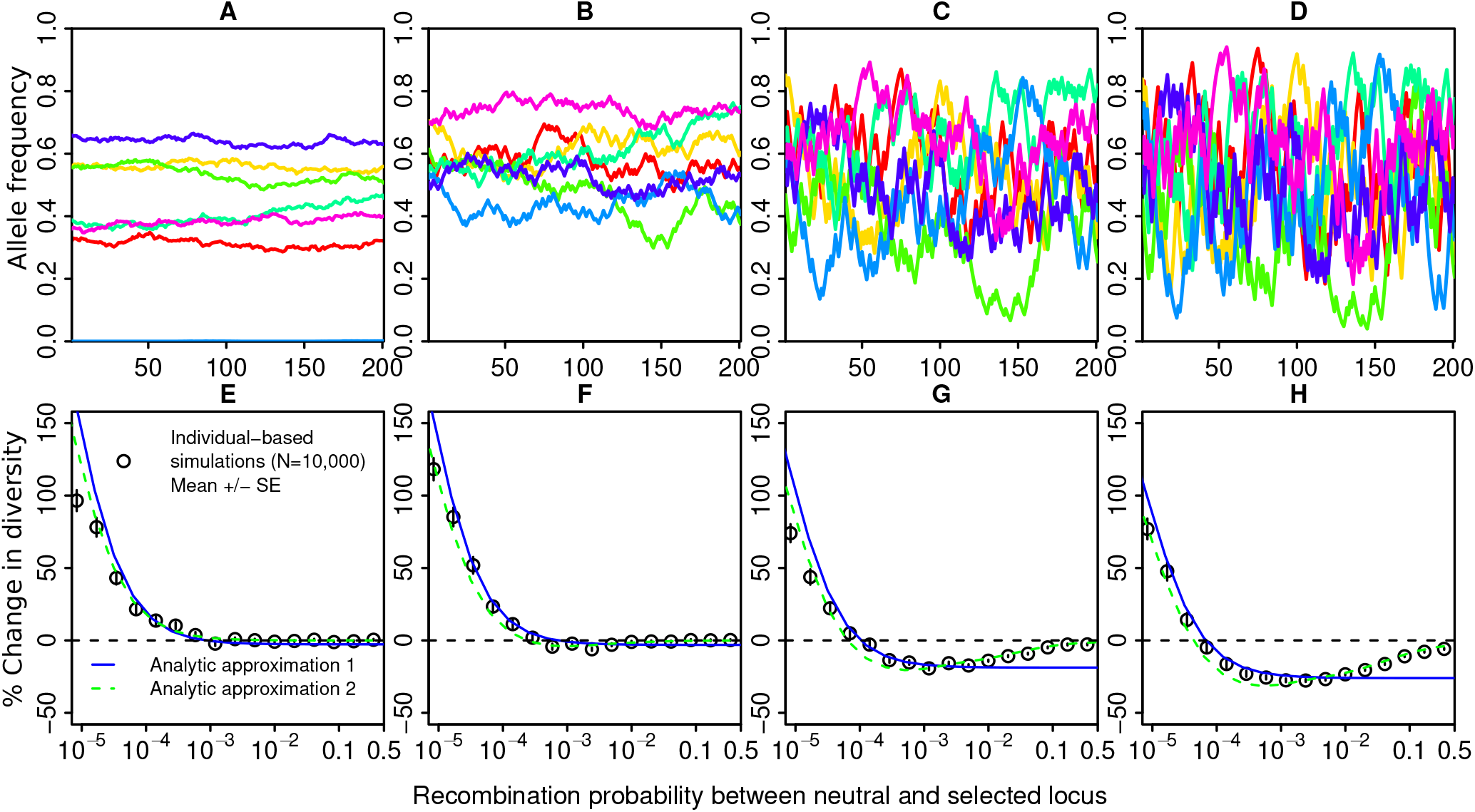
Results for the alternative scenario with uncorrelated selection across generations. The upper row shows seven replicate allele-frequency trajectories with A) *s* = 0.01, B) *s* = 0.1, C) *s* = 0.5, D) *s* = 1. The lower row depicts the corresponding genetic footprints. The simulation results are averages over 100 replicates. Other parameters: *h* = 0.6, *N* = 10, 000, *u* = 10^-6^.

### Chromosome-wide effect

Given that fluctuating balancing selection can lead to increases in diversity close to the selected site but decreases further away (see Figs. 3 and 4), we also quantify the net effect across the entire chromosome. Additional parameters in this analysis are *r*_0_, the per-generation recombination probability between adjacent base pairs, and *r_max_*, the recombination probability between the selected locus and the end of the chromosome. We assume that recombination occurs uniformly along the chromosome.

We quantify the net chromosomal effect in two ways. First, we run individual-based simulations with 40,000 sites between recombination distances *r*_0_ = 10^-6^ and *r_max_* = 0.49, with the same base-pair distance between all pairs of adjacent sites. We then average heterozygosity levels across this entire chromosome, including the selected site. Second, we use our analytical approximation (including both steps) to quantify the average heterozygosity for 1,000 recombination probabilities, *r*, between *r*_0_ and *r_max_*, evenly spaced on a logarithmic scale. Haldane’s map (Haldane 1919) then gives the corresponding distances in base pairs, *l*, to the selected site as

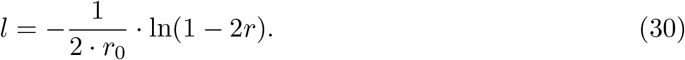

We then used the trapezoid rule to average over the footprint. For both approaches, we again relate average heterozygosity to the expected heterozygosity under neutrality.

There is good qualitative agreement between stochastic simulations and the analytical approximation (Fig. 6). Both suggest that the net chromosomal effect of fluctuating balancing selection is almost always negative. As expected from the results of the previous section, the reduction in average diversity levels is strong for fluctuations with a large amplitude (long seasons, large selection coefficients). For very weak selection, the analytical approximation predicts a very slight increase in diversity beyond neutral levels. However, the approximation ignores drift fluctuations in the allele-frequency trajectories, which become relevant for small s. Note that the error bars for the stochastic simulations overlap with zero, so we cannot distinguish whether the (very weak) average effect is positive or negative in this case. As with the local genetic footprint, the net chromosomal effect of fluctuating balancing selection intensifies over time, but with our parameter combinations already saturated after about 4N generations (Fig. A12).

**Figure 6.**
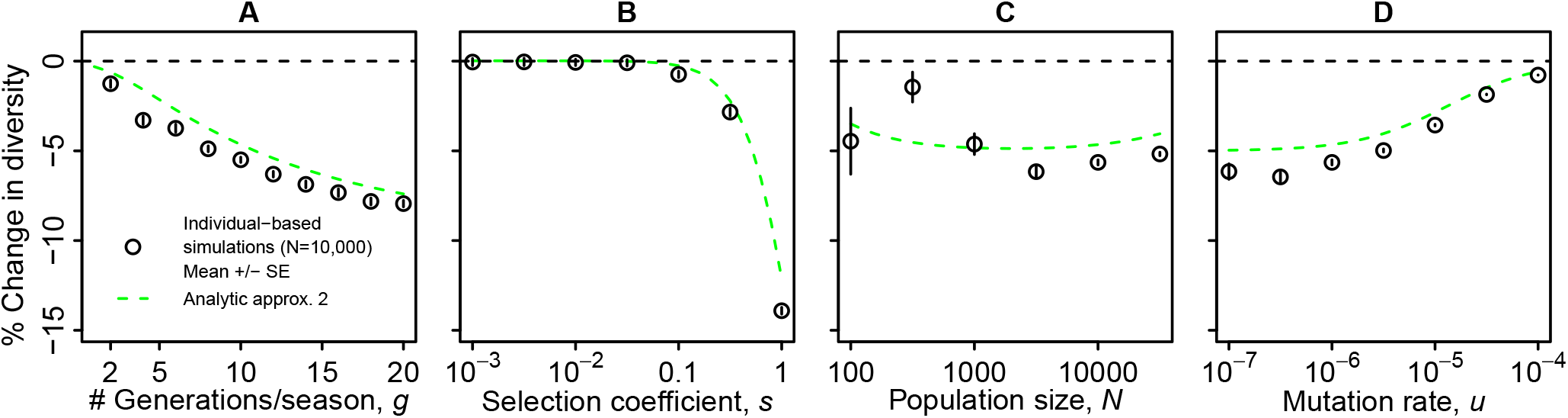
Comparison of analytical approximation 2 and stochastic simulations for the net effect of fluctuating balancing selection. Default parameter values: *r*_0_ = 10^-6^, *r_max_* = 0.49, *N_e_* = 10, 000, *u* = 10^-6^, *s* = 0.5, *h* = 0.6, *g* = 10. Averages over 40 replicates per data point with the last 10 sampling points taken into account. Vertical bars indicate standard errors. See Fig. A12 for the development of the net effect over time.

### Genome-wide effect of fluctuating balancing selection

We have seen that fluctuating balancing selection can impact diversity levels over large chromosomal regions and even in unlinked sections of the genome. Given these long-distance effects and given that many loci in the genome can be under fluctuating selection (e.g. on the order of hundreds in *Drosophila*, Bergland *et al*. 2014), the joint impact of multi-locus fluctuations may be substantial, even if the effect of each single fluctuating polymorphism is small. To quantify this impact, we randomly placed *n_l_* selected loci on a roughly *Drosophila*,-sized genome with three chromosomes of 50 Mb each. We used our individual-based simulation approach to simulate patterns of genetic diversity on the three chromosomes (see Appendix 8 for detailed methods). For simplicity, we focus on scenarios where all loci have the same parameters (s, h, g, and u), and assume multiplicative fitness (no epistasis) with per-locus contributions as shown in Table 1. In addition, we also ran simulations assuming that all selected loci are unlinked from each other (i.e. on different chromosomes) and unlinked from the focal neutral region, where the diversity is assessed.

Results for a range of selection coefficients, s, and numbers of selected loci, *n_l_* are displayed in Fig. 7. For sufficiently large selection coefficients and numbers of selected loci, we observed substantial genome-wide reductions in diversity. For the largest selection coefficient in this genome-wide simulation study (*s* = 0.1), allele-frequencies change on average by about 0.12 over one season and genome-wide diversity is reduced by about 30 %. Thus, the selection coefficients used here and the associated allele-frequency fluctuations are still relatively small (see Fig. A14) compared to those in Fig. 1 C and D underlying the pronounced local reductions in diversity in Fig. 3 C and D. As with the net chromosomal effect, the genome-wide effect took some time to build up, but was fully formed after about 4N generations (Fig. A15). Additional simulations, with selection coefficients drawn from an exponential distribution rather than being the same for all loci, yielded very similar results (Fig. A16).

**Figure 7.**
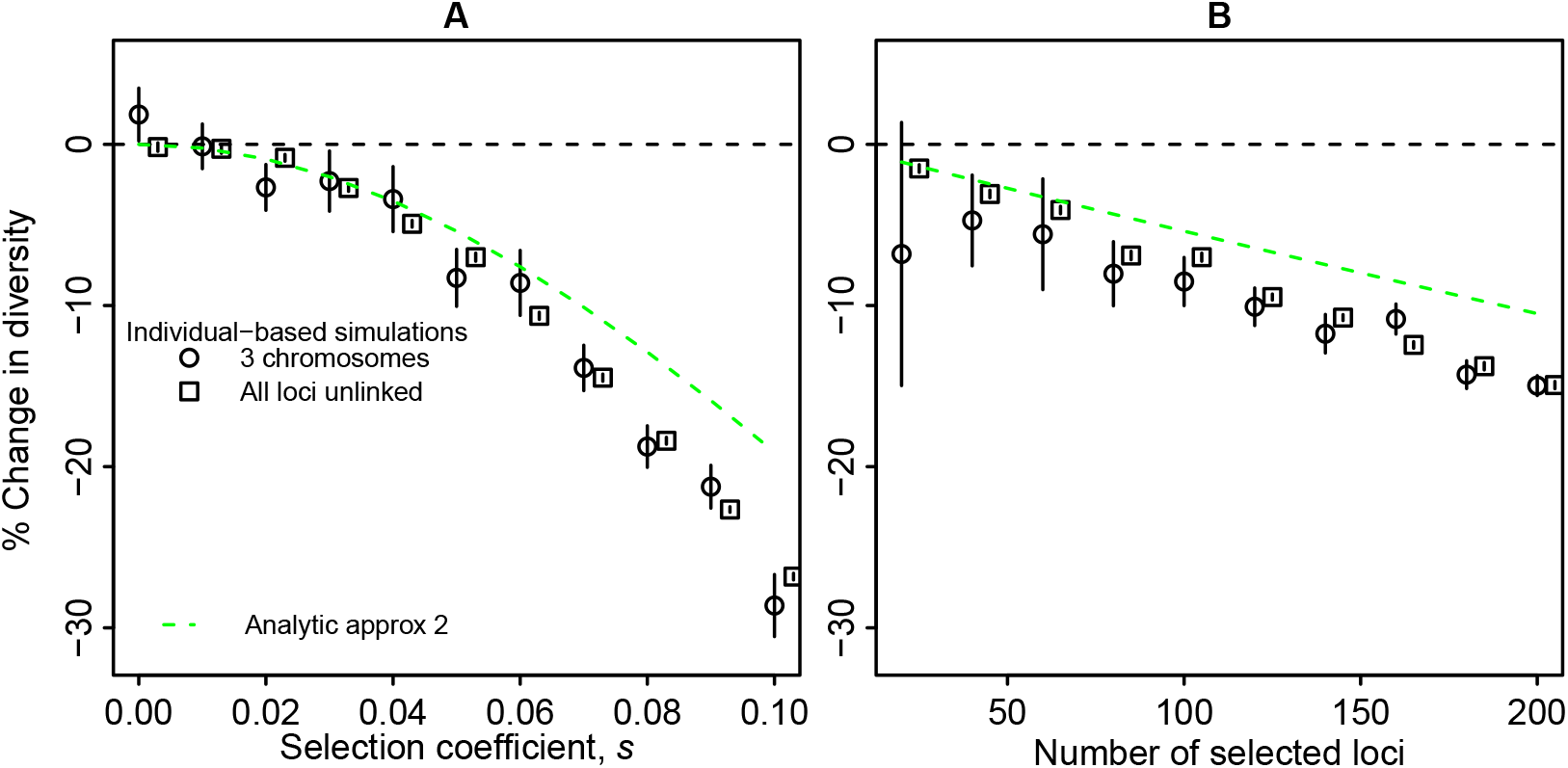
Relative change in genome-wide diversity due to fluctuating balancing selection at A) 100 randomly placed loci as a function of the selection coefficient, or B) different numbers of selected loci with *s* = 0.05. Each panel shows results for simulations where loci are randomly distributed over three chromosomes (circles, mean over 10 replicates ± standard error) and for corresponding simulations with all loci unlinked from each other and from the focal neutral region (squares, mean over 20 replicates ± standard error, slightly shifted to the right to avoid overlap). In each case, means are taken over the last 10 sampling points. Other parameters: *r*_0_ = 10^-6^, *N_e_* = 10, 000, *u* = 10^-6^, *h* = 0.6, *g* = 10. See Figs. A14 and A15 for information on the corresponding allele-frequency fluctuations and on the development of the footprint over time.

For the genome-wide footprint of balancing selection, it mattered little whether the 100 loci were distributed across three chromosomes or were each on a different chromosome, i.e. unlinked (Fig. 7 A). Thus, most of the genome-wide effect in this multi-locus model appears to be due to the effects of fluctuating balancing selection on diversity at unlinked loci (at *r* = 0.5).

In other words, the average region in the genome might be more affected by the cumulative long-distance effects of all loci in the genome that are under fluctuating balancing selection than by the short- or intermediate-distance effects of the nearest locus under selection.

We also compare the simulation results to a multi-locus extension of our full two-step analytical approximation. Here, we assumed a recombination probability of 0.5 for each locus and that every selected locus reduces the effective population size by the same factor. That is, we computed *F* from (22) for *r* = 0.5 and then used (14) to obtain

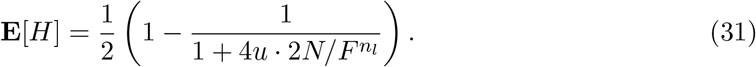

This approximation (green dashed line in Fig. 7) again somewhat underestimates the reduction in diversity due to fluctuating balancing selection, but predicts the qualitative shape of the curves. Notably, the effect of the selection coefficient is stronger than linear: strongly selected polymorphisms (with larger fluctuation amplitudes) matter disproportionately. In contrast, diversity levels decrease almost linearly with the number of loci under selection. Here, diversity levels for the smallest number of loci is well-predicted by (31), whereas for larger numbers of loci the actual diversity levels are somewhat lower than predicted based on the multiplicative reduction in effective population size across loci.

## Discussion

Our results indicate that fluctuating balancing selection leaves a characteristic footprint in linked neutral diversity, with a peak of genetic diversity close to the selected site, surrounded by diversity valleys in the flanking regions that extend to larger recombination distances. Although the increase at the peak is typically larger in magnitude, diversity reduction in the valleys affects much larger genomic regions, such that the net effect on chromosomewide diversity is almost always negative. Moreover, if multiple loci are under such fluctuating balancing selection, genome-wide diversity can be reduced substantially, even if fluctuations and footprints at individual loci are weak.

Using simulations and analytical approximations based on the structured coalescent and hitchhiking theory, we disentangled how the genetic footprint of fluctuating balancing selection is shaped by the various model parameters (see Fig. 4). We found that the depth of the diversity valley is determined by the harmonic mean allele frequency, 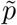, and thus mainly by the amplitude of the fluctuations, which, in turn, depends both on the selection strength per generation and the number of generations per season. Intuitively, with large fluctuations, lineages in the same background are forced through recurrent bottlenecks, which reduces diversity. While the width of the diversity peak is also governed by 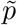 and becomes smaller for larger fluctuation amplitudes, the width of the diversity valley (i.e., its outer edge at large re-combination distances) only depends on the length of the season. It becomes wider for a large fluctuation frequency (small period). For a clear footprint to emerge, the population needs to be sufficiently large and selection sufficiently strong and sustained over an extended period of time. Although the dominance coefficient hardly influences the footprint, large enough dominance is required in our selection model to generate fluctuating balancing selection in the first place.

### Scope and limits of our approach

We developed an analytical approximation that consists of two steps. The first step accounts for the genetic structure of the population that matters most at small recombination distances. It relies on the structured coalescent. The second step accounts for the hitchhiking effect due to the fluctuations at larger distances, using results from Barton (2000). We show that both steps can be integrated to obtain an approximation that generally provides an excellent fit for the expected diversity levels across the entire recombination range. In our analytical approximation, the footprint is fully determined by the allele-frequency trajectory of the selected alleles. This has important consequences for the generality of our results.

First, we note that our results are expected to hold (qualitatively or even quantitatively) for a wide range of selection scenarios that lead to sustained allele-frequency fluctuations. In our model, stable fluctuations arise due to a specific form of selection with marginal overdominance. There are other mechanisms that can also lead to both stable polymorphism and allele-frequency fluctuations, such as temporal heterogeneity with a storage effect (see e.g. Chesson and Warner 1981; Gulisija *et al*. 2016; Park and Kim 2019; Reinhold 2000) or relative nonlinearity (e.g. Armstrong and McGehee 1980; Chesson 2000). However, as long as the resulting allele-frequency trajectories are comparable, the same analytical framework can be used, and the same footprint results. However, models that make very different assumptions, and for example include spatial structure or epistatic interactions (among multiple loci under fluctuating selection), will give rise to different footprints. Characterizing the footprints of balancing selection arising from various combinations of spatially and temporally heterogeneous selection (Svardal *et al*. 2015) remains an interesting question for future research.

Second, there are limits to the approach that lead to deviations in certain parameter regions. By using allele frequencies, rather than genotype frequencies, our approximation ignores that even within a background there is variance in reproductive success due to fitness differences between homozygotes and heterozygotes. This can lead to a slight overestimation of the chromosome-wide and genome-wide diversity levels. For multiple selected loci, we also ignore interference due to linkage disequilibria, which even occurs for unlinked loci. The fact that diversity decreases faster with increasing number of loci under fluctuating balancing selection than predicted by the simple multiplicative extension of our analytical approximation (see Fig. 7 B) could be explained by this effect.

It is currently still unclear whether the reduction in diversity at loosely linked or unlinked loci can be captured in terms of a single effective population size. The effects of strong purifying selection in a large population can at least be approximated by a reduction in effective population size and this approximation improves if the effective population size is made time-dependent (Nicolaisen and Desai 2012, 2013). Research on related complex forms of selection (Charlesworth and Jensen 2021; Taylor 2013), however, suggests that it might not always be possible to capture the effects of selection in terms of a single effective population size, but more work is needed to explore this in depth for fluctuating balancing selection.

For a closed analytical expression, we rely on a deterministic approximation for the allelefrequency trajectory. While this is appropriate for strong selection, it does not cover stochastic fluctuations due to drift, which become relevant for small selection coefficients or small population sizes (see Fig. A4 and A5). This leads to deviations of the analytical predictions from the simulation results, in particular very close to the selected site, where the analytical approach overestimates the diversity levels.

More general selection scenarios can be treated in a semi-analytical framework, where we condition the structured coalescent on a simulated, stochastic allele-frequency trajectory. In particular, this approach can also be used for selection in a random environment, as explored in our (limited) simulations for this case, with selection switching randomly from generation to generation (see Fig. 5). As in the seasonal case, the maximal reduction of the diversity (the bottom of the diversity valley) is again predicted by the harmonic mean frequency. However, the exact shape of recovery of diversity at large recombination rates differs from the case of seasonal selection, supposedly because the periods of the random fluctuations are not fixed, and typically larger for fluctuations with larger amplitude.

### Connections to prior research

Our general conclusion that genome-wide diversity levels can be substantially reduced is in accordance with previous results by Taylor (2013), Gillespie (1997) (both for the case of random fluctuations), as well as Park and Kim (2019) and Barton (2000). Taylor (2013) explored a structured-coalescent model for a locus under weak fluctuating selection without temporal autocorrelation where the allele-frequency dynamics can be approximated by a diffusion process. Despite the very different selection regime with frequent allelic turnover, his numerical results also show an excess of variation at or very close to the selected site itself and a decrease in diversity at more loosely linked sites. In stochastic simulations, Gillespie (1997) also found a long-range reduction in diversity in simulations of some parameter combinations of the SAS-CFF (stochastic additive scale - concave fitness function) model of temporally variable selection, a footprint that was however hard to distinguish from that of other models based on Tajima’s D. Interestingly, here effective population size decreased less than linearly with population size, whereas our analytical predictions suggest a linear increase with population size, at least when conditioned on a certain allele-frequency trajectory.

Park and Kim (2019) considered a haploid population inhabiting two patches, where one experiences selection with a cyclically fluctuating optimum and the other acts as a refuge, such that polymorphism can be maintained via a storage effect. Despite the quite different biological scenario, they also found that levels of polymorphism at linked neutral loci can be either higher or lower than under neutrality. If there is a reduction, they attribute this also to the recurrent sweep-like patterns at the selected loci. Finally, Barton (2000), provides a quite general treatment linking the theory of hitchhiking to fluctuating selection quantifying the reduction in effective population size as a function of recombination distance from the selected loci, results that we heavily relied on for the second step of our analytical approximation.

A relevant empirical study is the evolve-and-resequence experiment by Huang *et al*. (2014). Here, the authors compared *Drosophila melanogaster* populations evolving in four different environments: a constant salt-enriched or cadmium-enriched environment, a spatially heterogeneous, and a temporally variable environment. Although the study could not directly identify the actual targets of selection, they used proxies to classify sites as closely linked or further away from likely targets of selection. They found that for closely linked sites, populations in the spatially heterogeneous environments had the highest diversity, followed by the temporally variable environment, with the constant environments having the lowest diversity. Interestingly, for sites further away, populations in the spatially heterogeneous environment still had the highest diversity, but now the constant environments were intermediate, and the temporally variable environments had the lowest levels of genetic diversity, which is consistent with our predictions.

### Outlook: Implications for genome scans and genome-wide diversity levels

As mentioned in the Introduction, one of the motivations for characterizing the expected footprints of various types of selection is to be able to scan genomes for specific types of selection. Although our results suggest a characteristic “peak-and-valley” pattern, it may often be difficult to detect in real data. While the diversity peak close to the selected site is often narrow and high, and thus conspicuous, the valley is typically shallow and broad. The footprint will then be difficult to distinguish from “simple” balancing selection with constant frequency of the polymorphism at the selected locus. Only for large allele-frequency fluctuations, a clear footprint with a detectable peak and valley arises. In this case, the shape of the footprint can look similar to the “volcano” pattern generated by adaptive introgression (Setter *et al*. 2020). However, the width of the diversity valley is typically much broader for fluctuating balancing selection (recombination distances up to about 1/number of generations per cycle) than for the introgression volcano, where the width is governed by the selection strength. Going beyond diversity levels, future work will need to see whether the site-frequency spectrum (SFS) or haplotype patterns can provide additional clues for the detection of fluctuating balancing selection. Results by Huerta-Sanchez *et al*. (2008) suggest that fluctuating selection can under some conditions leave signatures in the SFS that can be distinguished from other types of selection. Alternatively, recent developments in detecting selection from data sets with multiple sampling points (Buffalo and Coop 2020) could be fruitful also for detecting fluctuating balancing selection.

Even if fluctuations are small and local footprints might be hard to detect, the net effect on the chromosome and, if multiple loci are under fluctuating balancing selection, on the genome can be substantial. It has been known for a long time that genome-wide levels of supposedly neutral genetic variation do not correlate well with census population sizes, contrary to what one would expect under neutral evolution (Lewontin’s paradox, Buffalo 2021; Charlesworth and Jensen 2022; Leffler *et al*. 2012; Lewontin 1974; Maynard Smith and Haigh 1974). Levels of genetic diversity in large populations are often much smaller than predicted based on census sizes. Although demographic factors probably also play a large role (Charlesworth and Jensen 2022; Ellegren and Galtier 2016), there is increasing evidence that the effects of linked selection at many loci in the genome are at least contributing to this pattern (Corbett-Detig *et al*. 2015; Elyashiv *et al*. 2016). So far, the main modes of selection that are thought to reduce diversity levels disproportionately in large populations are selective sweeps and background selection, i.e., the loss of variation due to the elimination of linked deleterious mutation (Charlesworth and Jensen 2021; Elyashiv *et al*. 2016). Our results now suggest that fluctuating balancing selection could be another mode of selection playing an important role in shaping levels of genome-wide genetic variation.

## Acknowledgements

This work was supported by a Lise-Meitner postdoctoral fellowship from the Austrian Science Foundation FWF to MJW (project number M 1839-B29). We would like to thank Nick Barton for an important hint and Christian Huber for helpful comments on an earlier version of the manuscript.

## Online Appendix

## Appendix 1 Analytical approximation of allele-frequency trajectories

Focusing on symmetric scenarios and neglecting the effect of mutation on allele frequencies, we can state a recursion for the ratio of the two allele frequencies. For *i* < *g*, the winter allele is favored and we have

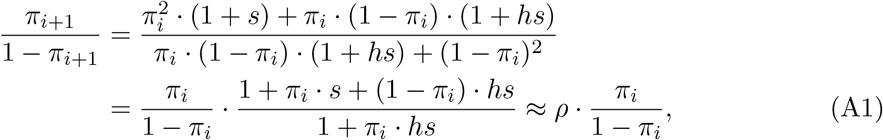

where 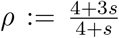. This approximation is based on the assumption that both *h* and *π_i_* are close to 0.5. Thus, *π*/(1 – *π*) grows by a factor *ρ* every generation. We could also keep h as a parameter in the growth factor rather than setting it to 0.5, but it turned out that this does not add much accuracy.

Solving this geometric growth model, the allele frequencies during winter (*i* ≤ *g*) are

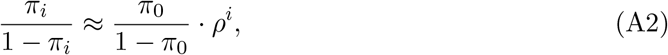

where *π*_0_ is the frequency of the winter allele at the beginning of the year, i.e. its minimum allele frequency along the cycle. To achieve symmetry, the two alleles must exchange their allele frequencies after g generations such that we have

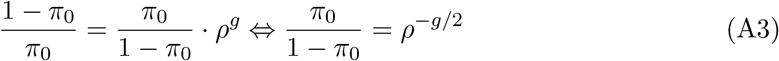

and thus

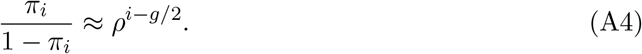

Hence the frequencies along the cycle are

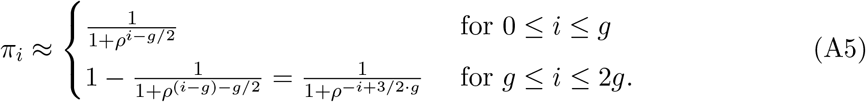

The above approximation is based on an assumption of constant, in particular frequency-independent, fitness effects over the course of one season. In symmetric cases, the gains in frequency over one season balance exactly the losses in the other season. In asymmetric situations these gains and losses are not generally balanced. Coexistence of alleles in those cases is only possible via negative frequency-dependent selection due to marginal overdominance. Thus an approximation based on constant (frequency-independent) fitness cannot work. Thus, for asymmetric cases we use numerically-determined cyclic allele-frequency trajectories as a basis.

## Appendix 2 Expected heterozygosity

Since we consider a symmetric, biallelic mutation model, there can be back-mutations, which becomes important if the product of expected coalescence time and mutation rate is large. Two sampled lineages with coalescence time *T* will carry different alleles if and only if there is an odd number of mutations along the branches of length 2*T* in their genealogy. The number of mutations is Poisson-distributed with parameter 2*uT*. Thus, the probability that the two sampled lineages carry the same allele is

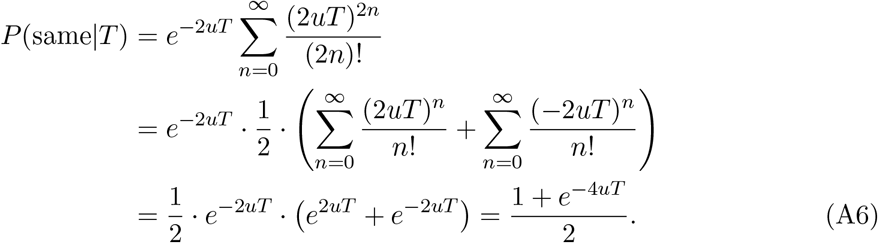

Assuming that coalescence time is approximately exponentially distributed with parameter λ, we obtain:

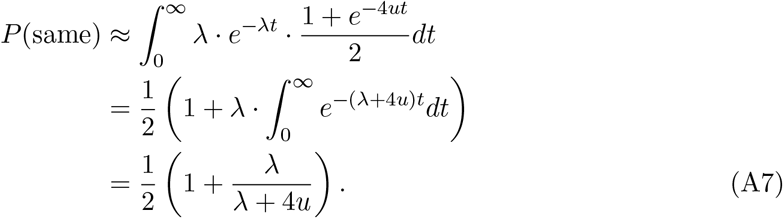

Then for sample configuration (*i,j*), we use λ = 1/**E**[*T*_(i;j)_]. Defining the expected heterozygosity *H* = 1 – *P*(same), i.e. as the probability that the two lineages carry different alleles, we obtain

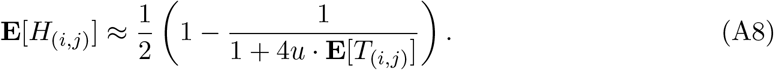

That is, as the product of mutation rate and expected coalescence time increases, heterozygosity increases, but with diminishing returns and a saturation at 0.5 in our biallelic model (see Fig. A1). Using this formula instead of the classical 2*u***E**[*T*]/(1 + 2*u***E**[*T*]) improved the fit between simulations and analytical results. Under neutrality, **E**[*T*_(i,j)_] = 2*N* and (A8) coincides with the result in Ewens (2004), p. 97.

**Figure A1.**
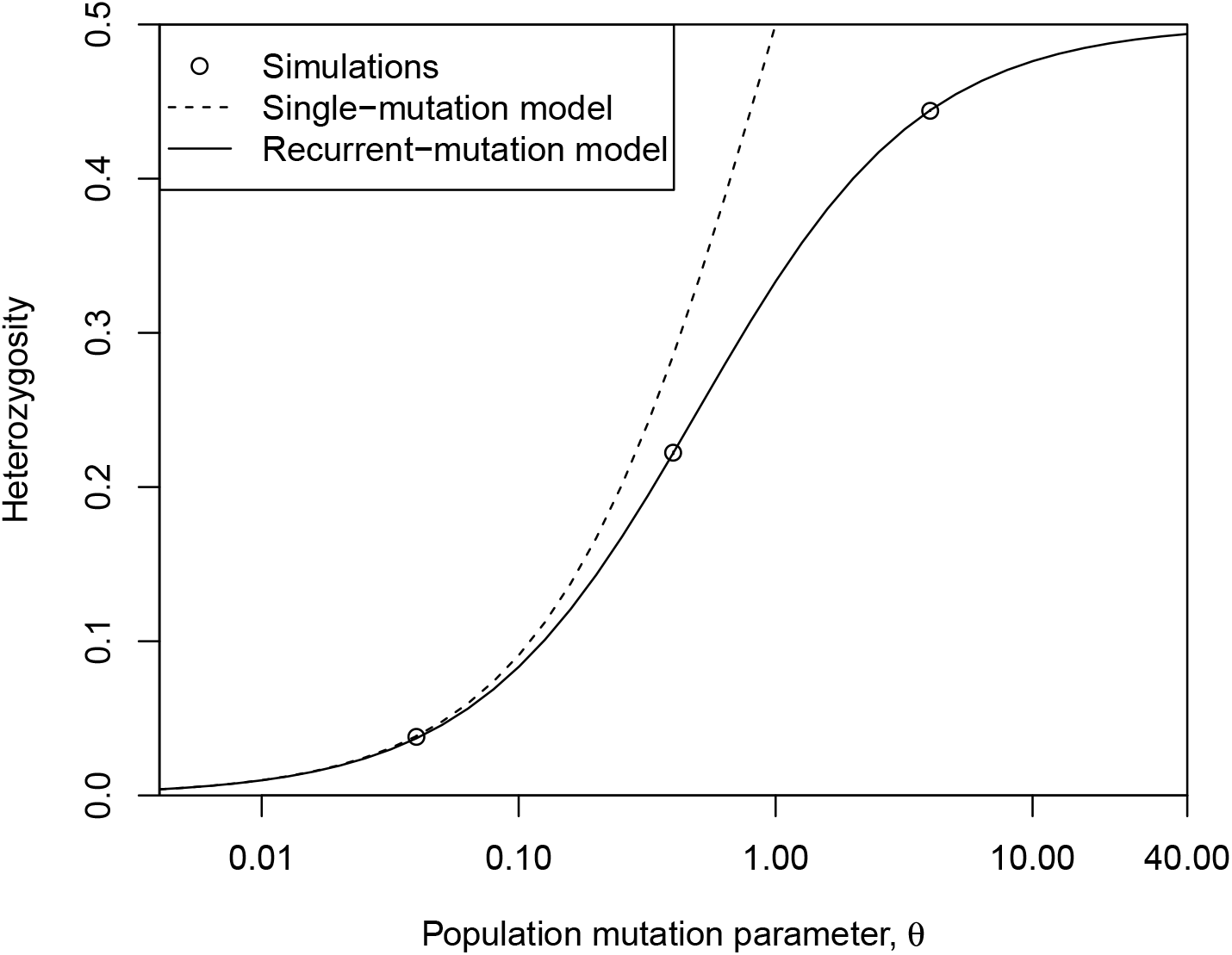
Expected heterozygosity under neutrality as a function of the population mutation parameter, *θ* = 4*Nu*. Simulation results are for *N* = 10, 000 and each point represents the mean over 16 replicates. The corresponding standard errors are too small to be visible. The single-mutation model uses the standard formula (2 · *u* · **E**[*T*])/(1 + 2 · *u* · **E**[*T*]) whereas the multi-mutation model is based on (14). Both use **E**[*T*] = 2*N*.

## Appendix 3 Convergence of step 2 of the analytic approximation for small recombination rates

To understand how this approximation connects to the analytical approximation in (11) and (12), we derive the limit of *F* as the recombination rate *r* goes to zero. Using the shorthand 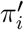 for 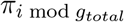, we have for *S_i_* (26) for small *r*:

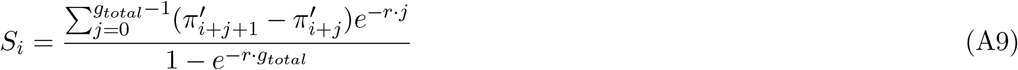

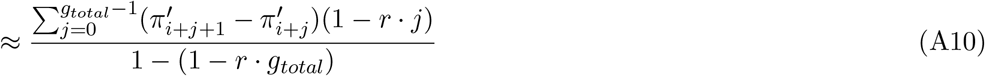

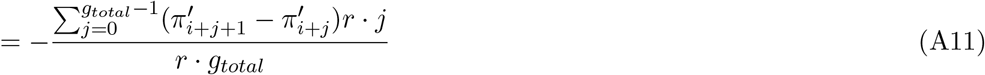

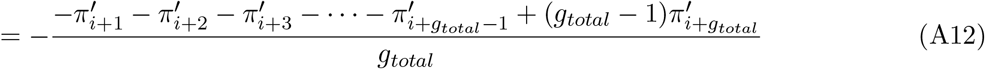

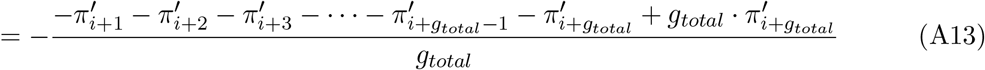

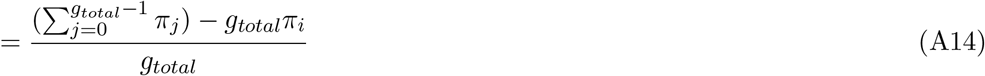

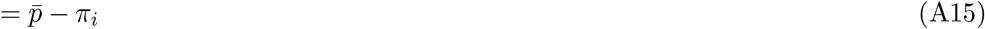

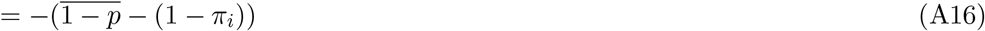

Inserting this in (22) gives:

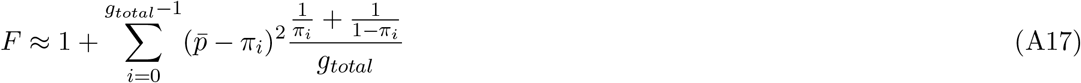

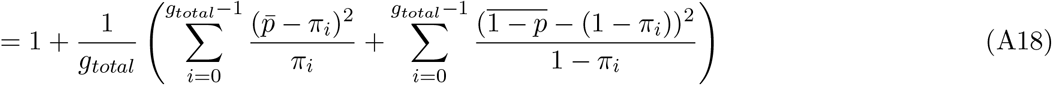

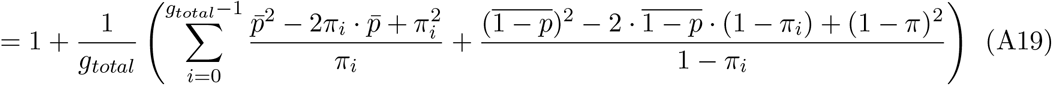

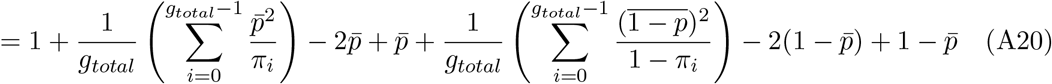

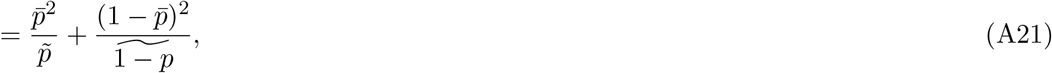

where~denotes the harmonic mean over a cycle.

In symmetric cases, 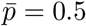 and 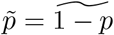 and we have for small *r*

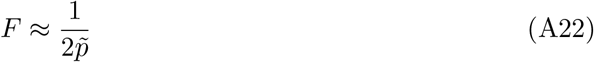

and

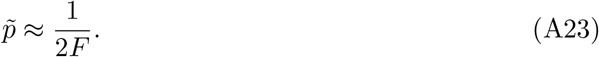

## Appendix 4 Initialization of individual-based simulations

Under neutrality, the number of copies M of an arbitrarily chosen allele evolves according to the following model:

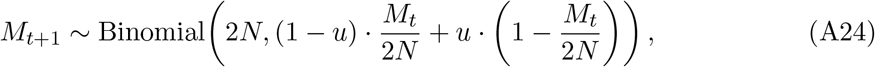

where *N* is the population size and *u* is the mutation probability. To initialize the initial allele frequencies in the population, we start at allele frequency 0.5 and iterate (A24) for 0.1*N* · (*L* + 10) generations, where L is the number of sites that we want to simulate. Every 0.1N generations, we take the current allele frequency (which can be 0 or 1) to initialize the frequency at one of the sites (throwing away the first 10 to allow for some burn-in. We randomly shuffle the vector of allele frequencies. simuPOP is then randomly assigning genotypes to individuals in the initial populations such that these allele frequencies are respected.

## Appendix 5 Additional results for the genetic footprint

**Figure A2.**
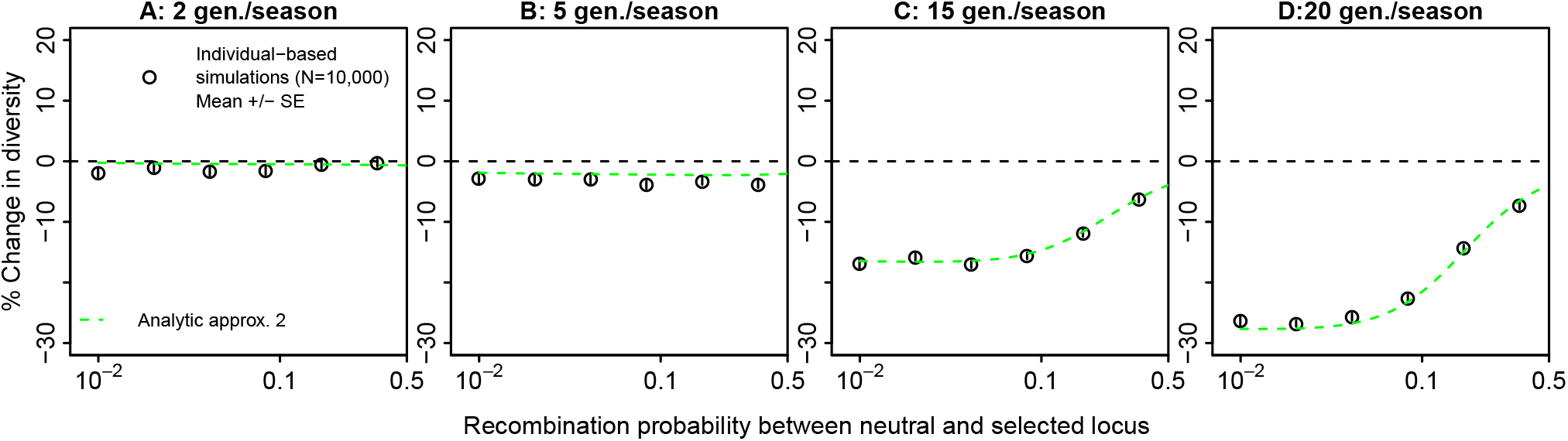
Zoom into the region of large recombination rate in Fig. 3.

**Figure A3.**
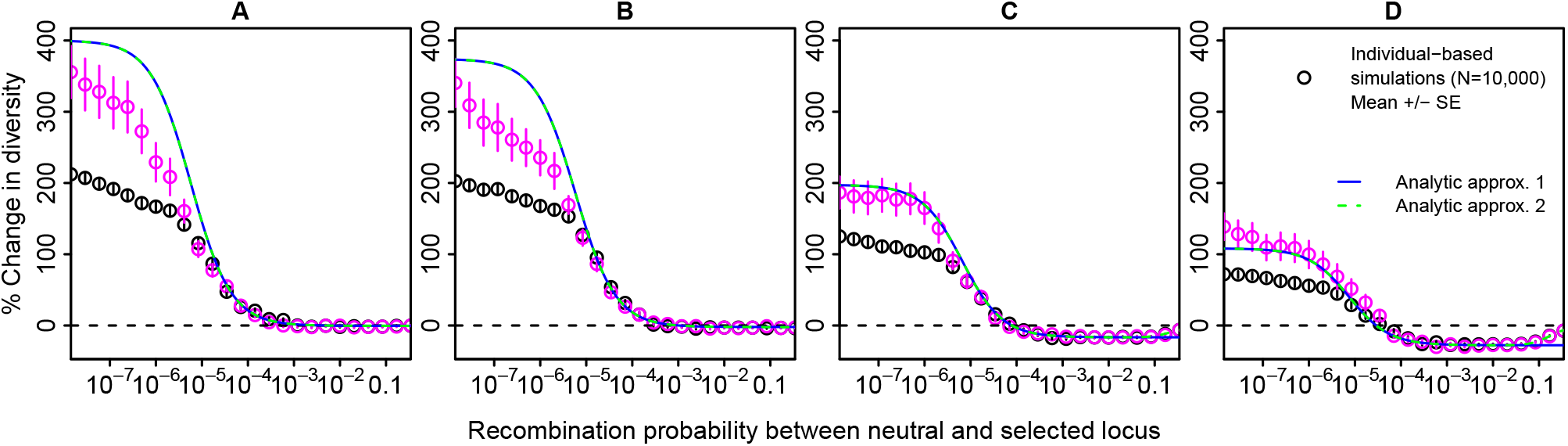
Genetic footprint of fluctuating balancing selection for various season length, *g*, as in Fig. 3, but now including the region very close to the selected site with a low recombination probability between neutral and selected locus. A) *g* = 2, B) *g* = 5, C) *g* = 15, and D) *g* = 20. The y-axis represents the relative change in diversity compared to the neutral case. Black points are averages over 100 replicates with five sampling points each. Pink points are averages over 20 replicates run for 450 instead of 15 *N* generations with 29 sampling points 0.5 *N* generations apart at the end of the simulation run.

In Fig. 3, we found a discrepancy between analytical approaches and simulation results close to the selected site. Such a discrepancy can arise if diversity levels have not fully equilibrated by the end of the simulation. The large diversity levels close to the selected site are due to very large coalescence time and our simulation might not have been long enough to cover the tail end of the distribution, thus leading to a slight underestimation of diversity levels close to the selected site. This is consistent with the fact that diversity levels were still increasing in the final phase of the simulation close to the selected site in Fig. A9 and with the finding in Fig. A3 that the simulated diversity levels increased and more closely matched the analytical approximation when simulations were run for longer.

**Figure A4.**
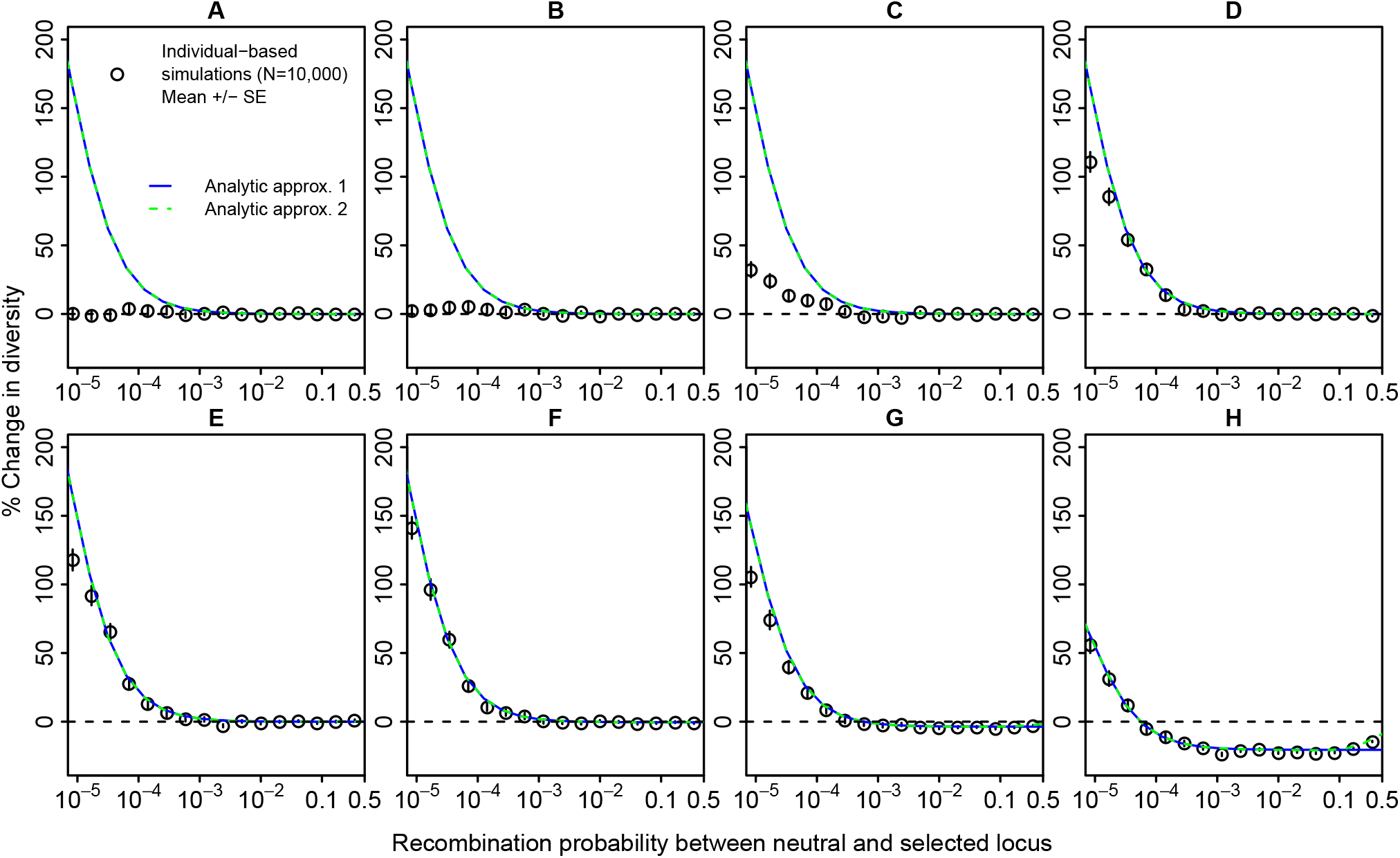
Genetic footprints of fluctuating balancing selection for various selection coefficients, *s*. In A) *s* = 0, and the values of *s* in B) to H) are 10^-3^,10^-2^’^5^,10^-2^,10^-1^’^5^,10^-1^,10^-0^’^5^,1. The y-axis represents the relative change in diversity compared to the neutral case. Points are averages over 100 replicates with five sampling points each. Note that the stochastic simulations in A) are truly neutral, whereas the analytical approximations with s = 0 erroneously assume constant allele frequencies of 0.5, i.e. constant balancing selection.

**Figure A5.**
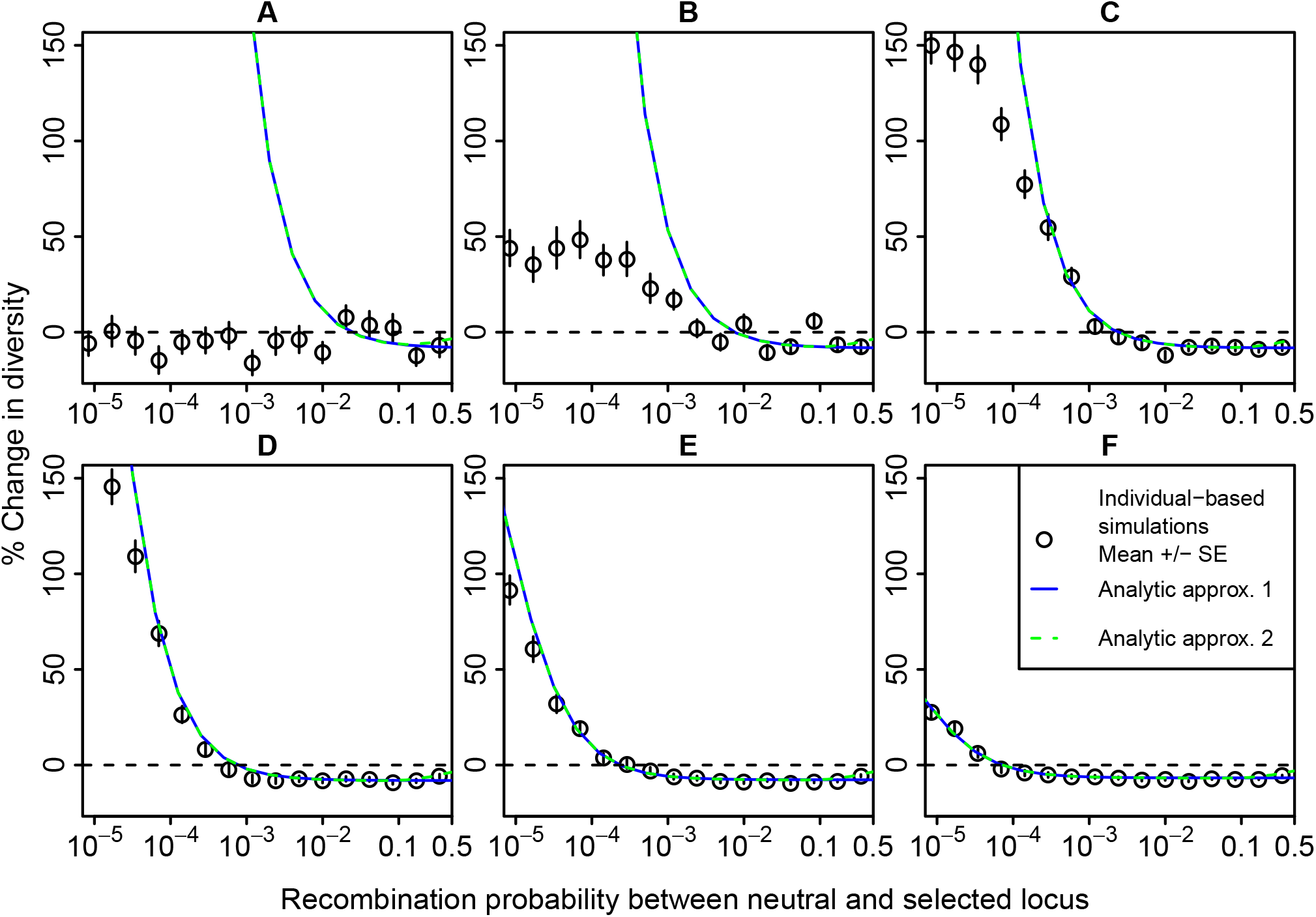
Genetic footprints of fluctuating balancing selection for various population sizes, *N*. A) *N* = 100, B) *N* = 316 = ⌊10^2.5^⌋, C) *N* = 1, 000, D) *N* = 3,162 = ⌊10^3.5^⌋, E) *N* = 10, 000, F) *N* = 31, 623 = ⌊10^4.5^⌋, where the “floor” function ⌊*x*⌋ gives the largest integer smaller or equal to *x*. The y-axis represents the relative change in diversity compared to the neutral case. Points are averages over 100 replicates with five sampling points each.

A striking effect of increasing the mutation rate u is that both the increase in diversity close to the selected site and the decrease further away decrease in magnitude (Fig. A6). This is the case because in our biallelic and symmetric mutation model, heterozygosities start to saturate at high mutation rates and are then less sensitive to fluctuating balancing selection (see ι Appendix 2). However, this effect appears to be somewhat specific to our biallelic symmetric mutation model. In reality, there will be four bases available at each site and mutation rates are often much smaller such that diversity levels may be far away from saturation.

**Figure A6.**
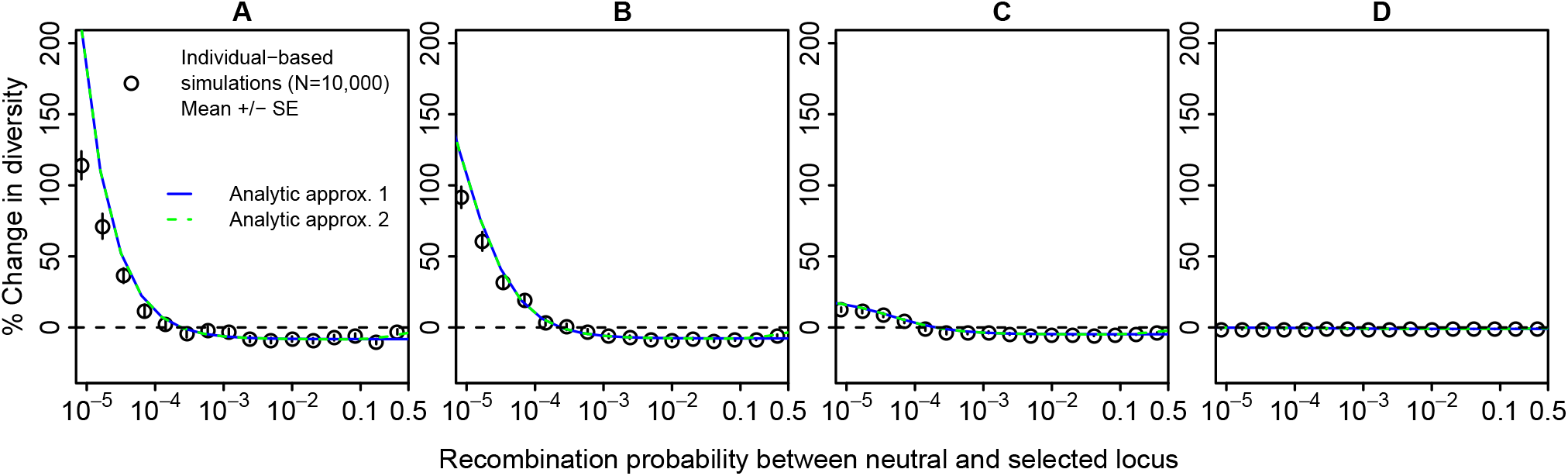
Genetic footprints of fluctuating balancing selection for various mutation rates, *u*. A) *u* = 10^-7^, B) *u* = 10^-6^, *u* = 10^-5^, *u* = 10^-4^. The y-axis represents the relative change in diversity compared to the neutral case. Points are averages over 100 replicates with five sampling points each.

**Figure A7.**
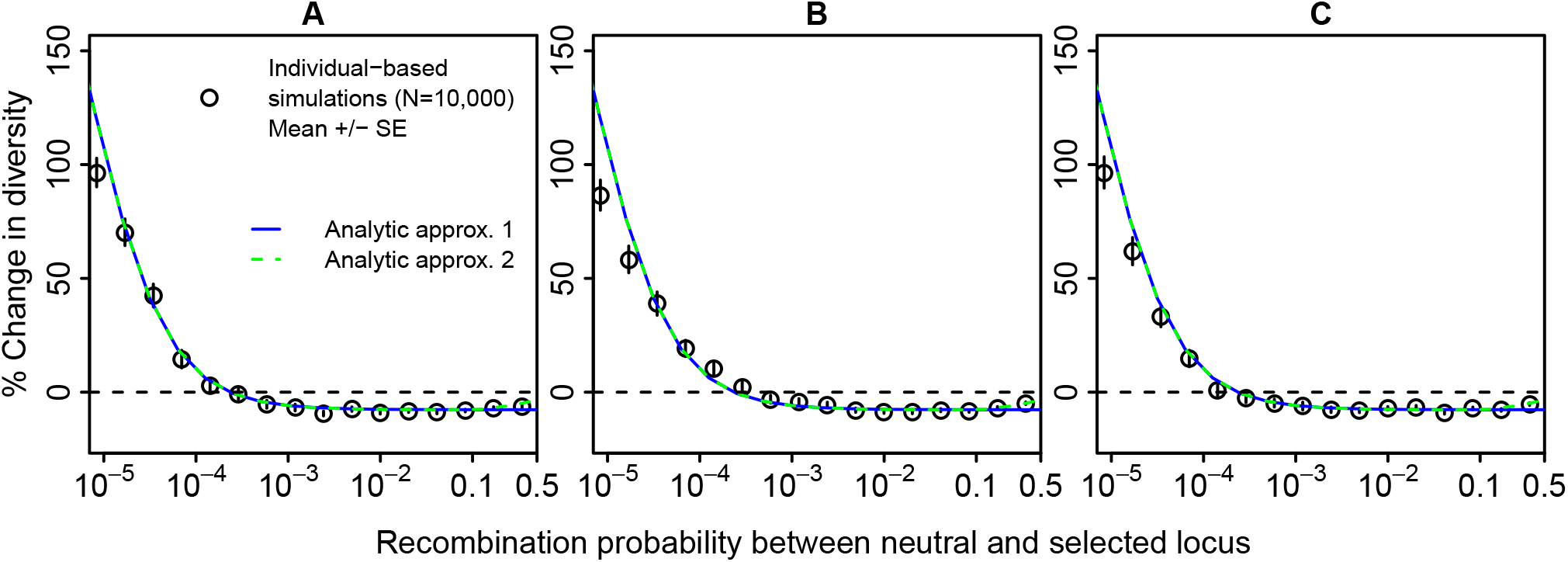
Genetic footprints of fluctuating balancing selection for various dominance coefficients, *h*. A) *h* = 0.5, B) *h* = 0.75, and C) *h* =1. The y-axis represents the relative change in diversity compared to the neutral case. Points are averages over 100 replicates with five sampling points each.

**Figure A8.**
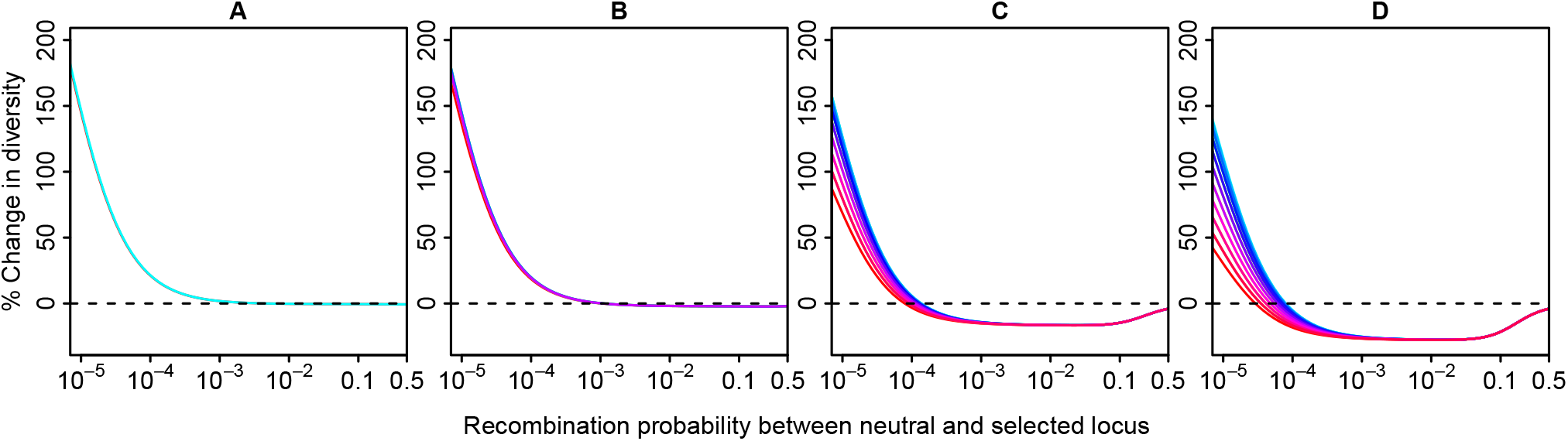
The expected local footprint of selection changes depending on the generation of the seasonal cycle in which the sample is taken. Curves based on analytic approximation 2. Red curves correspond to sampling times when allele frequencies are rather extreme whereas shades of blue represent sampling points where allele frequencies are more intermediate.

**Figure A9.**
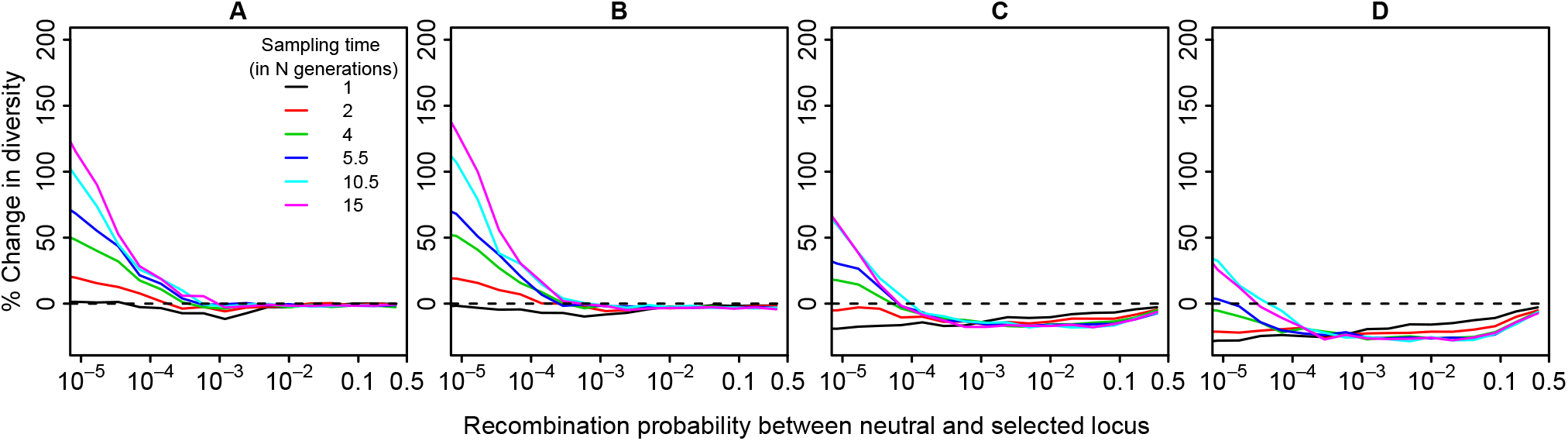
Development of genetic footprints over time for the scenarios in Fig. 3, i.e. with A) *g* = 2, B) *g* = 5, C) *g* = 15, and D) *g* = 20. The curves represent the average over 100 replicates of relative change in diversity compared to the neutral case at different sampling times. Sample i is taken (*i* + 1)/2 · *N* generations after the onset of the seasonal selection pressure.

## Appendix 6 Asymmetric scenarios

We assume that

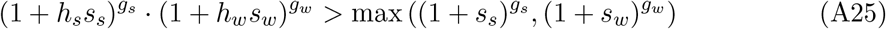

such that there is marginal overdominance and thus protected polymorphism. Since our analytical approximation for allele-frequency trajectories (Appendix 1) does not work for asymmetric scenarios, we determine the cyclical allele frequency trajectories numerically by iterating (1) and (2).

The structured coalescent Markov process in asymmetric cases is determined by four parameters: 1) the coalescent rate within the winter-allele background 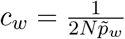, where 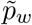 is the harmonic mean frequency of the winter allele, 2) the coalescent rate within the summerallele background 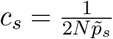, where 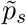 is the harmonic mean frequency of the summer allele, 3) the rate at which lineages move from the winter background to the summer background 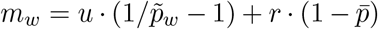, where 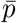 is the arithmetic mean winter allele frequency, and 4) the rate at which lineages move from the summer background to the winter background 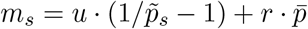.

With these transition rates, the recursions for the expected coalescence times for the three possible starting configurations are

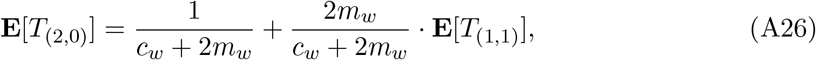

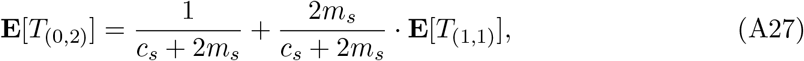

and

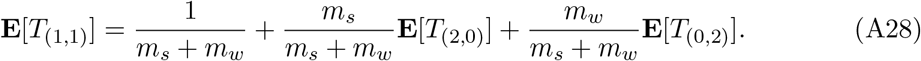

Solving for **E**[*T*_(2,0)_], **E**[*T*_(0,2)_], and **E**[*T*_(1,1)_] yields

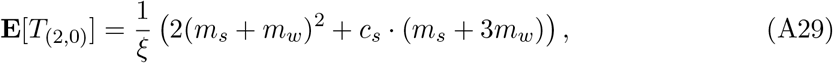

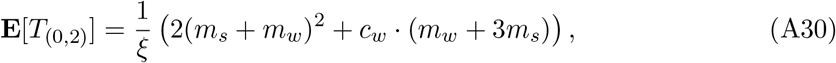

and

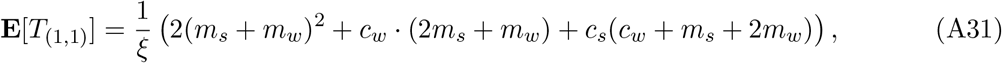

where

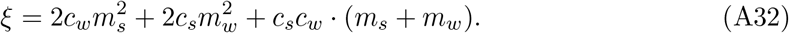

Note that

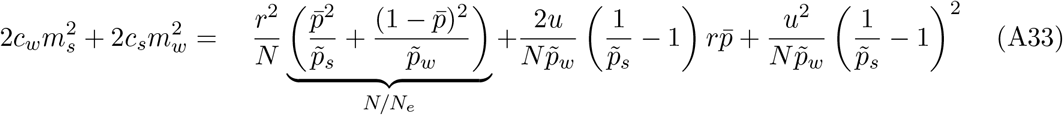

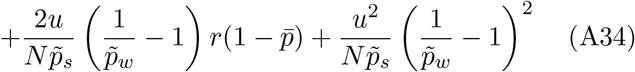

To take the second step of the analytical approximation, we then note that the term with the under-brace corresponds to *N/N_e_* (see (27)) and is also the limit of *F* as *r* → 0 in the general asymmetric case (see Appendix 3). We can thus replace the term with the underbrace by *F*. Thus we set

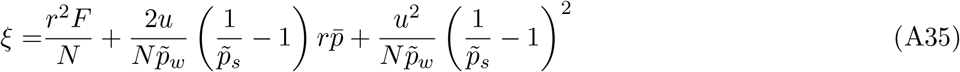

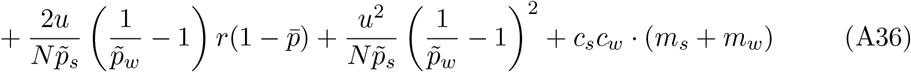

in (A29)–(A31) to complete the second step of the analytic approximation in the asymmetric case.

Again, the expected coalescence times are then transformed to relative changes in heterozygosity as explained above for the symmetric case. Fig. A10 shows genetic footprints of fluctuating selection for different degrees of asymmetry.

**Figure A10.**
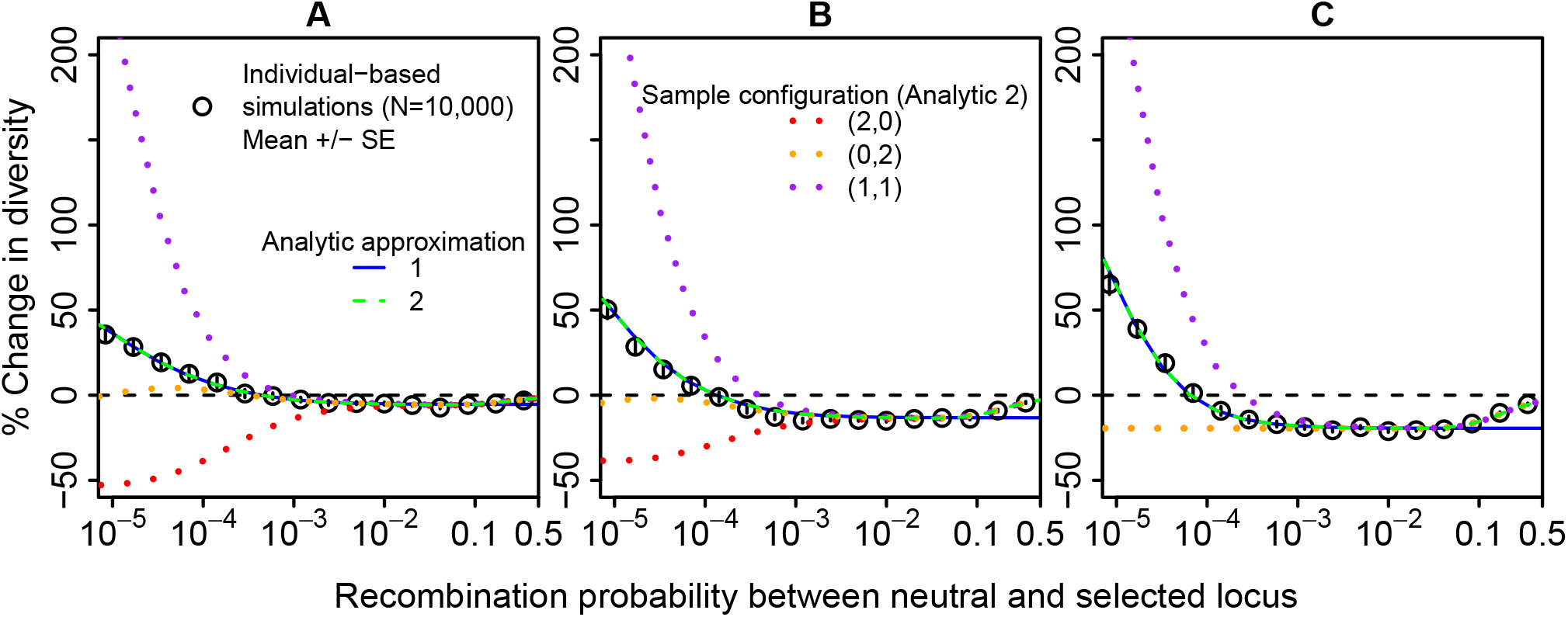
Genetic footprint of fluctuating balancing selection with different degrees of asymmetry in the number of generations per season. (A) *g_s_* = 33, *g_w_* = 7, (B) *g_s_* = 28, *g_w_* = 12, (C) *g_s_* = *g_w_* = 20. That is, the total number of generations per cycle is 40 in all cases. Points are averages over 100 replicates with five sampling points each. The analytic approximations are shown for a random sample from the population. In addition analytic approximation 2 is shown for the different sample configurations. Other parameters: *s_s_* = = 0.5, *h_s_* = *h_w_* = 1,*N* = 10, 000, *u* = 10^-6^. The corresponding allele-frequency trajectories are shown in Fig. A11.

**Figure A11.**
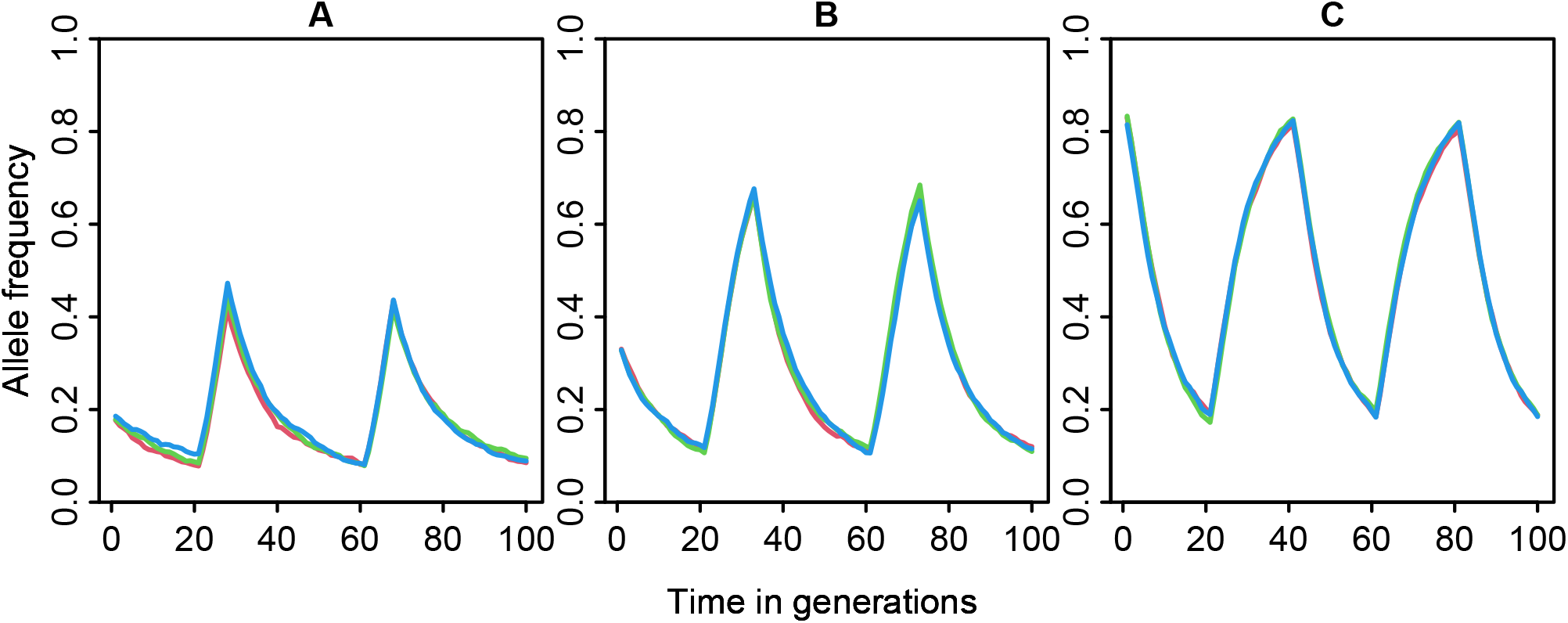
Allele-frequency trajectories of the winter allele with different degrees of asymmetry in the number of generations per season. (A) *g_s_* = 33, *g_w_* = 7, (B) *g_s_* = 28, *g_w_* = 12, (C) *g_s_* = *g_w_* = 20. Other parameters: *s_s_* = *s_w_* = 0.5, *h_s_* = *h_w_* = 1,*N* = 10, 000, *u* = 10^-6^.

## Appendix 7 Additional results for the chromosome-wide effect

**Figure A12.**
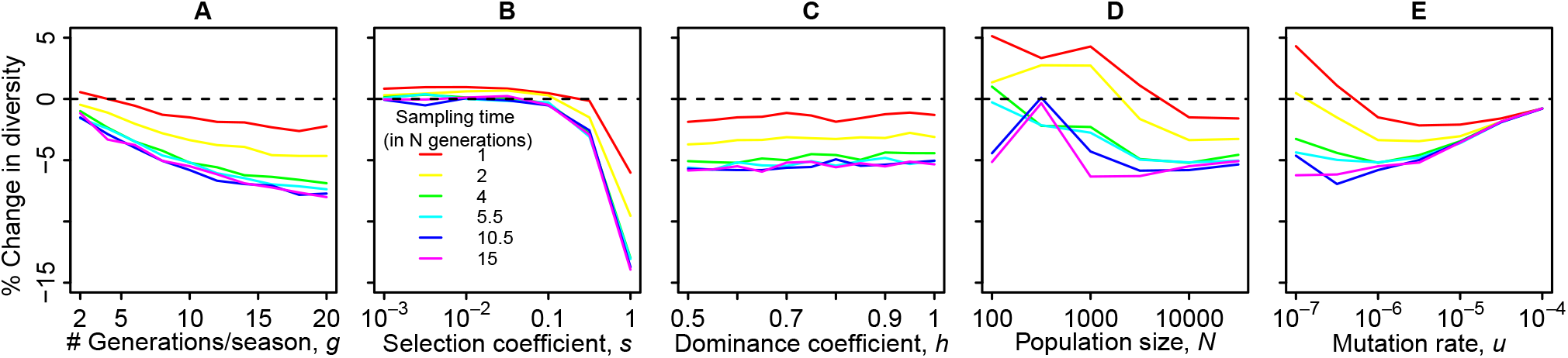
Development over time of the net effect of fluctuating balancing selection on chromosome-wide diversity in the individual-based simulations. Samples were taken every 0.5 *N* generations, with the first sample 0.5 *N* generations after the onset of fluctuating balancing selection. Default parameter values: *r*_0_ = 10^-6^, *r_max_* = 0.49, *N_e_* = 10, 000, *u* = 10^-6^, *s* = 0.5, *h* = 0.6, *g* = 10. 40 replicates per data point. Note that the increase in diversity in early samples with small population sizes and small mutation rates appears to be an artifact because the burn-in time underlying the initial allele-frequency distribution (see Appendix 4) was a bit too short for these parameter combinations.

## Appendix 8 Quantifying the effects of multiple randomly-placed selected loci on genome-wide diversity

The *n_l_* selected loci are placed randomly on the chromosomes, that is, the number of selected loci on each chromosome is drawn from a multinomial distribution, and then for each of the three chromosomes the positions of the respective numbers of loci under selection are drawn uniformly without replacement.

In addition to the selected loci, we simulated a number of neutral sites on each chromosome. To capture the rapidly-changing diversity landscape around the selected site without having to simulate too many sites, we selected the positions of the neutral sites according to the following scheme: We chose a minimum and maximum recombination probability *r_min_* and *r_max_* and set up a vector of recombination probabilities to a selected locus with *L_r_* elements such that the ith element is

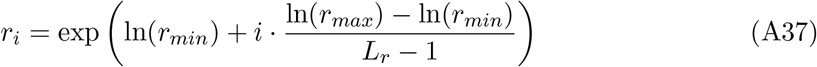

for *i* ∈ {0,1,..., *L_r_* – 1}. That is, we choose the recombination probabilities evenly spaced on a logarithmic scale. In all simulations shown, we use *L_r_* = 500.

For each selected locus, we simulate neutral loci at distances given by *r_i_* to the left and right, but only up to the midpoint of the stretch to the next selected locus. We place an additional neutral locus at the midpoint and then switch to the recombination vector for the next locus. We proceed analogously at the ends of the chromosome.

Let *ρ_i_* be the recombination probability between the *i* – 1th and the *i*th of these sites. To compute *ρ_i_*, we notice that a recombination event between the selected locus and locus *i* occurs if there is a recombination event between the selected locus and locus *i* – 1 or a recombination event between locus *i* – 1 and locus *i*, but not both. Thus

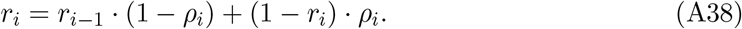

Solving for *ρ_i_* gives:

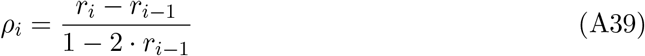

for *i* ∈ {1,2,..., *L_r_* – 1}. We set *r_min_* to *r*_0_ the assumed recombination probability between adjacent sites per generation. An example recombination map with just four loci is shown in Fig. A13. We again used Haldane’s map (30) and the trapezoid rule as described above for the net chromosomal effect to obtain average diversity levels across the chromosome. For further details of the implementation, please refer to the python and R scripts in the supplement.

As a comparison, we ran simulations with all selected loci on different chromosomes and a focal neutral region, also unlinked, with 40,000 sites evenly spaced on a chromosome with a recombination probability 0.49 between its two extremes.

**Figure A13.**
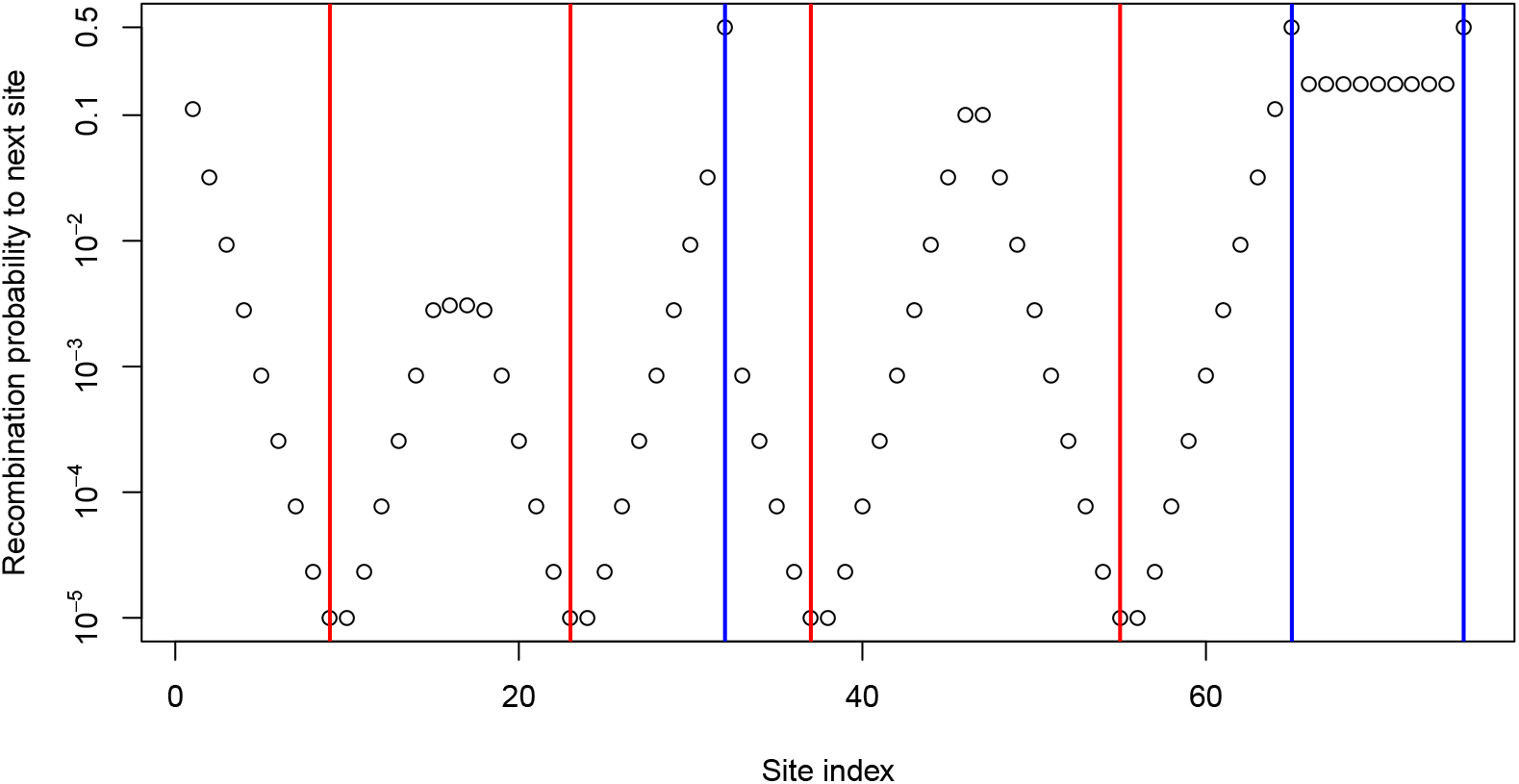
Example for a chromosome map with four selected loci thrown onto three chromosomes. Blue vertical lines represent the end of a chromosome. Red vertical lines represent loci under selection. For each site, the point represents the recombination probability to the next site to the right.

**Figure A14.**
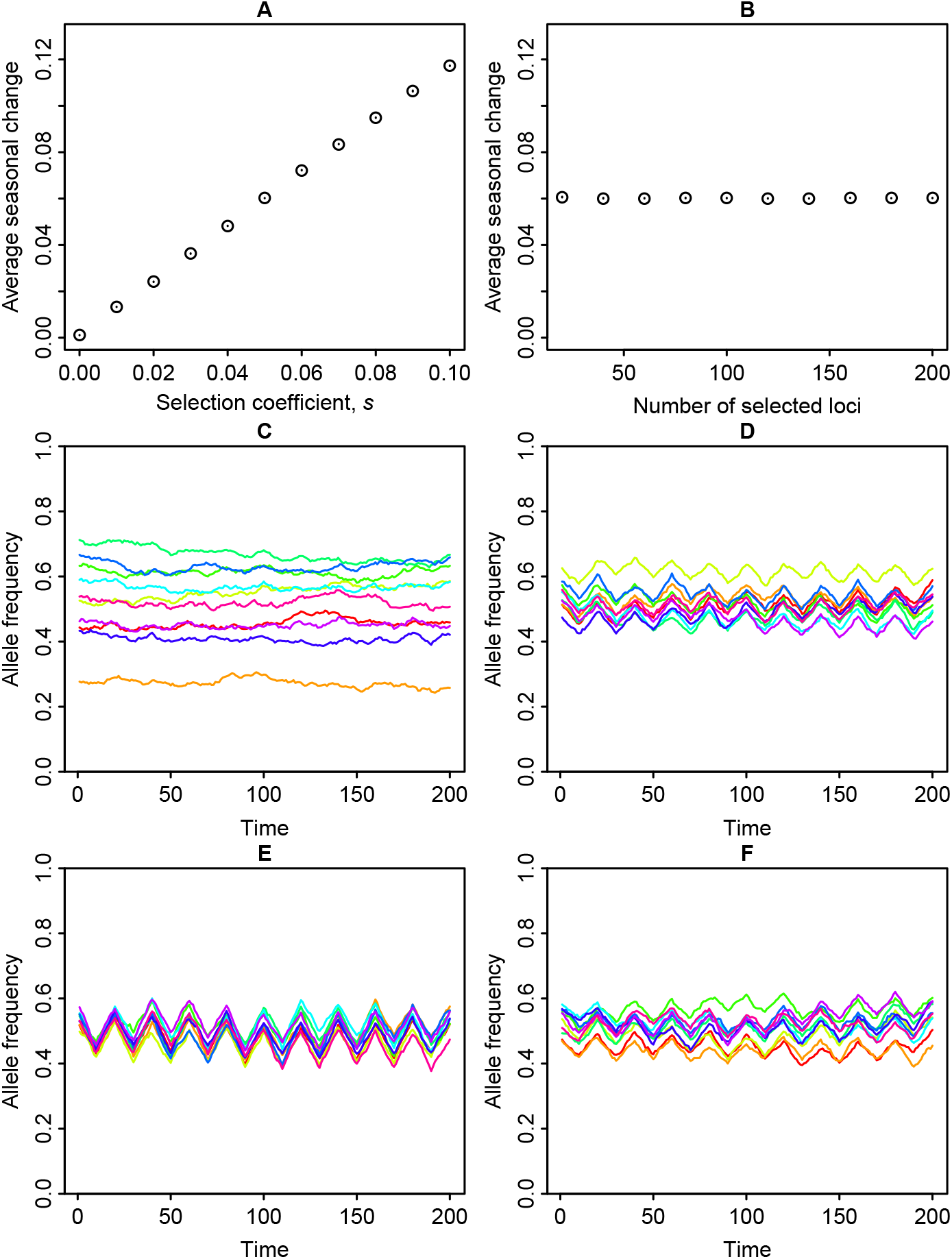
Allele-frequency dynamics corresponding to the 3-chromosome scenario in Fig. 7. Average absolute value of the allele-frequency change over one season as a function of (A) the selection coefficient, and (B) the number of selected loci, and example allele-frequency trajectories for 10 randomly selected loci (C-F). In (A, C, and E), there are 100 loci under selection. In (C), *s* = 0.02, and in (E), *s* = 0.1. In (B, D, and F), *s* = 0.05. The total number of selected loci is 40 in (D) and 200 in (F).

**Figure A15.**
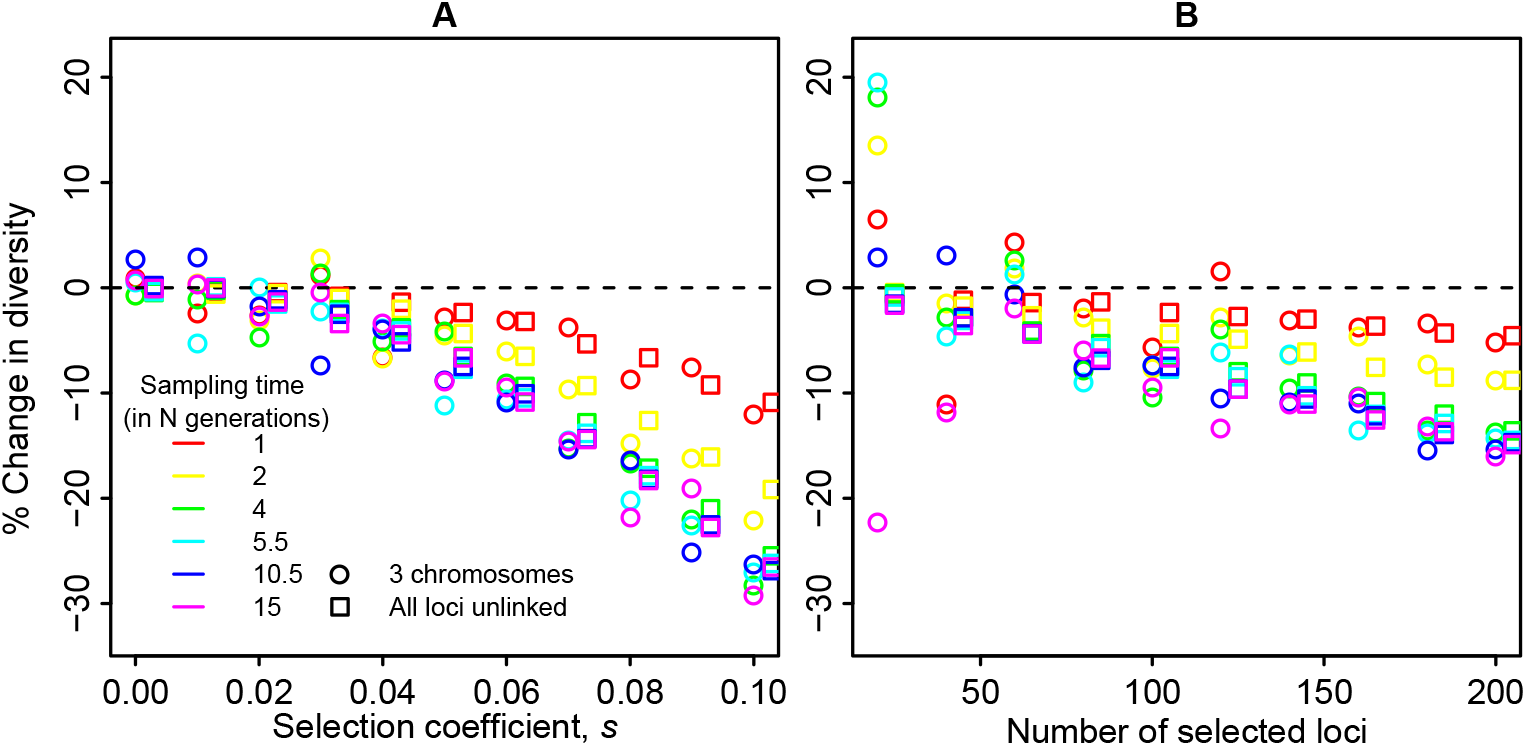
Temporal development of the impact of fluctuating balancing selection on genomewide diversity in the scenarios shown in Fig. 7. A) 100 randomly placed loci with different selection coefficients. B) Different numbers of selected loci with *s* = 0.05. Each panel shows results for simulations where loci are randomly distributed over three chromosomes (circles, mean over 10 replicates) and for corresponding simulations with all loci unlinked from each other and from the focal neutral region (squares, mean over 20 replicates, slightly shifted to the right to avoid overlap). Other parameters: *r*_0_ = 10^-6^, *N_e_* = 10, 000, *u* = 10^-6^, *h* = 0.6, *g* = 10.

**Figure A16.**
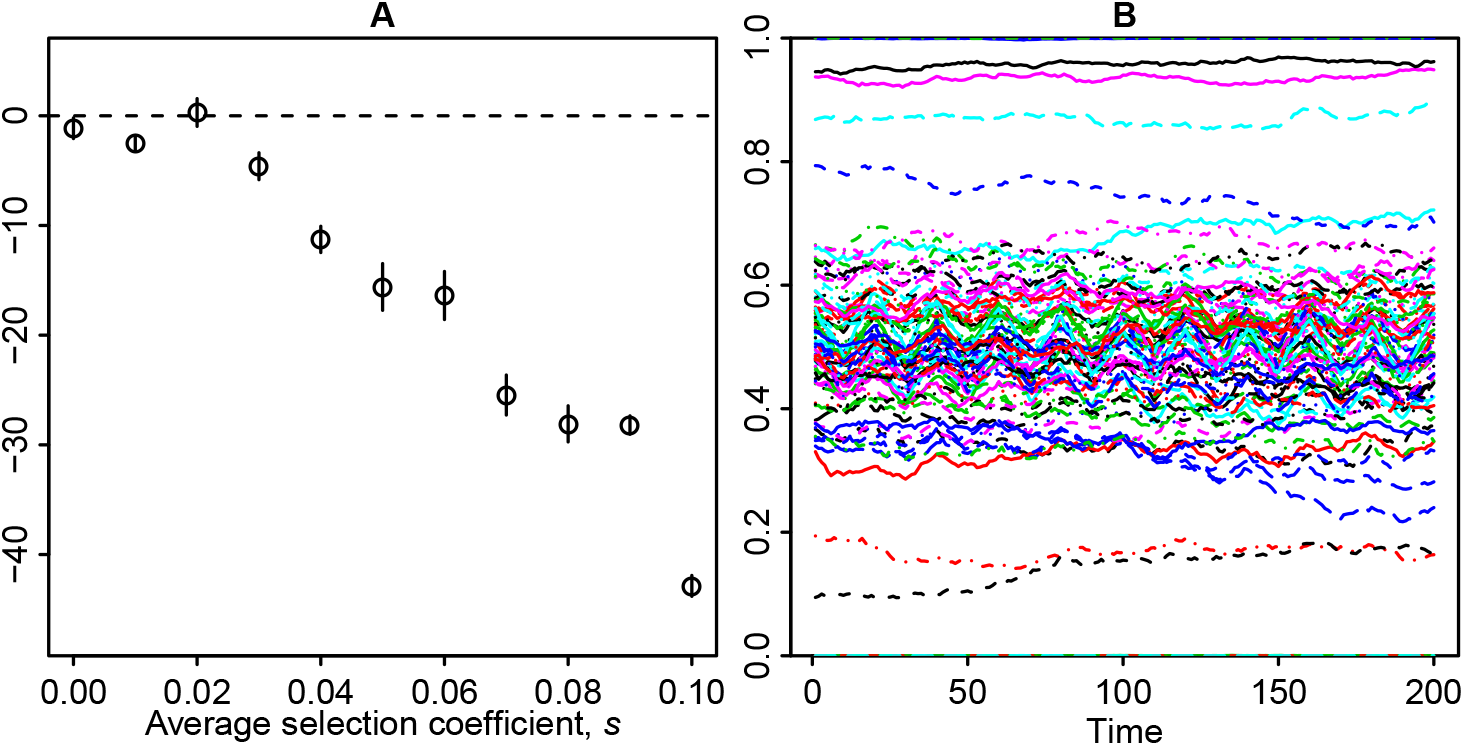
Fluctuating balancing selection at 100 randomly placed loci with exponentially distributed selection coefficients (with *s_w_* = *s_s_*). A) Relative change in genome-wide diversity (mean ± standard error over 10 replicates, sampling points 11 to 29) as a function of the average selection coefficient. Allele-frequency fluctuations at the 100 loci over the final 200 generations in one of the replicates. Other parameters: *r*_0_ = 10^-6^, *N_e_* = 10, 000, *u* = 10^-6^, *h* = 0.6, *g* = 10.

